# Bisphenol A induces sex-dependent alterations in the dynamics of neuroendocrine seasonal adaptation in Djungarian hamsters

**DOI:** 10.1101/2024.02.12.580037

**Authors:** Marie-Azélie Moralia, Béatrice Bothorel, Virginie Andry, Valérie Simonneaux

## Abstract

In nature, species synchronize reproduction and energy metabolism with seasons to optimize survival and growth. While the effects of endocrine-disrupting chemicals (EDCs) exposure on conventional laboratory rodents are increasingly studied, their impacts on mammalian seasonal adaptation remain unexplored. This study investigates the effect of oral exposure to bisphenol A (BPA) on physiological and neuroendocrine seasonal adaptation in Djungarian hamsters. Adult female and male hamsters were orally exposed to BPA (5, 50, or 500 µg/kg/d) or vehicle during a 10-week transition from a long (LP) to short (SP) photoperiod (winter transition) or vice versa (summer transition). Changes in body weight, food intake, and pelage color were monitored weekly and, at the end of the exposure, gene expression of hypothalamic markers of photoperiodic, reproductive and metabolic integration, reproductive organ activity, and glycemia were assessed. Our results revealed sex-specific effects of BPA on acquiring SP and LP phenotypes. During LP to SP transition, females exposed to 500 µg/kg/d BPA exhibited delayed body weight loss and reduced feed efficiency associated with a lower expression of *somatostatin* in the arcuate nucleus (ARC), while males exposed to 5 µg/kg/d BPA showed an accelerated acquisition of SP-induced metabolic parameters. During SP to LP transition, females exposed to 5 µg/kg/d BPA displayed a faster LP adaptation in reproductive and metabolic parameters, along with quicker ARC kisspeptin downregulation and delayed ARC *Pomc* upregulation, while males exposed to BPA exhibited decreased expression of central photoperiodic integrators without changes in the physiological LP acquisition. This pioneering study investigating EDC impacts on mammalian seasonal physiology shows that BPA alters the dynamic of metabolic adaptation to both SP and LP transitions with marked sex dimorphism, causing temporal discordance in seasonal adaptation between males and females. These findings emphasize the importance of investigating EDCs’ impact on non-conventional animal models, providing insights into wildlife physiology.

**Highlights:**

- Djungarian hamster’s seasonal adaptation is disrupted by BPA oral exposure
- BPA delays in females and accelerates in males the metabolic adaptation to short days
- BPA accelerates in females, not in males, metabolic/reproductive adaptation to long days
- BPA affects the photoperiodic expression of central reproductive and metabolic genes

Graphical abstract

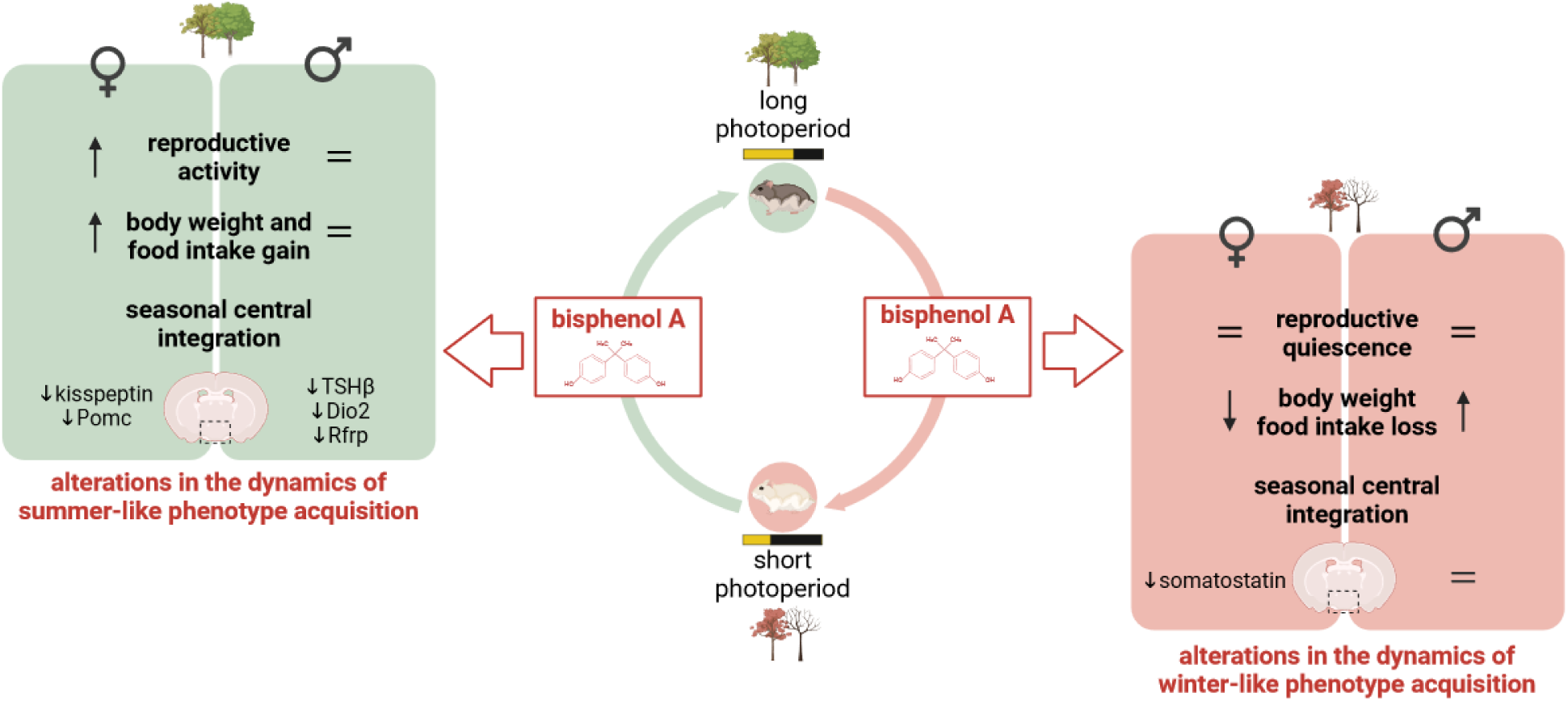

## 1. Introduction

Among the plethora of adaptations to environmental constraints that species are capable of, physiological adaptations to seasonal changes are particularly remarkable. Indeed, most species living in moderate to high-latitude regions of the world synchronize their metabolic and reproductive physiology to seasonal changes in geophysics factors (*i.e*. temperature, humidity, photoperiod), enabling them to optimize their energy resource allocation and the timing of the birth of their offspring throughout the year, ultimately guaranteeing the survival of the species (Norris and Jones, 1987; Bronson, 1985; Shinomiya et al., 2014). A typical example of such adaptive processes is observed in the Djungarian hamster (*Phodopus sungorus*), a seasonal rodent that exhibits high food consumption and body weight, active reproduction, and gray-colored fur in spring and summer, while, as winter is coming, it displays reduced metabolic activity, inhibited reproduction, and white fur (Bartness and Wade, 1984; Warner et al., 2010; Figala et al., 1973; Duncan and Goldman, 1984).

Decades of studies using seasonal species such as Djungarian and Syrian hamsters, and sheep unveiled the neuroendocrine mechanisms underlying these physiological adaptations (Dardente and Simonneaux, 2022; Hazlerigg and Simonneaux, 2015). Seasonal species use the annual change in the 24-hour day length, *i.e*. photoperiod, as a reliable cue to determine the seasons (Nakane and Yoshimura, 2019). In mammals, annual photoperiodic changes are integrated into the brain by a retino-hypothalamic-pineal axis resulting in a long synthesis of the nocturnal hormone melatonin during the long winter nights (or short photoperiod, SP) and in a short melatonin synthesis during the short summer nights (or long photoperiod, LP) (Hazlerigg and Simonneaux, 2015). This melatoninergic seasonal message is integrated into the *pars tuberalis*, an endocrine structure of the pituitary gland that contains a high density of melatoninergic receptors (Klosen et al., 2002; Dardente et al., 2003). There, thyrotropic cells express the thyroid hormone stimulating beta subunit (TSHβ) which is strongly inhibited by the long melatonin peak in SP (Bockmann et al., 1996; Dardente et al., 2003). In LP, the highly produced TSH activates TSH receptors localized on tanycytes, glial cells of the intracerebroventricular-hypothalamic interface, which in turn regulate the expression of thyroid hormone metabolism enzymes called deiodinases (Tu et al., 1997; Guadaño-Ferraz et al., 1997; Yoshimura et al., 2003). Thus, the LP-induced expression of TSH results in both a higher expression of deiodinases 2 (Dio2) that convert the prohormone thyroxine (T4) to the bioactive thyroid hormone triiodothyronine (T3), and a reduced expression of Dio3 that inactivates T3, whereas the SP-induced inhibition of TSH is associated with an opposite high expression of Dio3 and low expression of Dio2 (Watanabe et al., 2004; Revel et al., 2006; Hanon et al., 2010; Sàenz de Miera et al., 2013; Yoshimura et al., 2003; Helfer et al., 2013). This melatonin-driven TSH control of tanycytic Dios leads to a photoperiodic switch in intra-hypothalamic concentration of T3, with higher values in LP as compared to SP (Klosen et al., 2013). Through mechanisms yet to be determined, seasonal variations in hypothalamic T3 regulate the expression of the GnRH regulators kisspeptin in the arcuate nucleus (ARC) and RFRP-3 in the dorsomedial hypothalamus (Klosen et al., 2013; Henson et al., 2013, Dardente and Simonneaux, 2022), as well as the ARC growth hormone regulator somatostatin (Herwig et al., 2012, 2013; Petri et al., 2014; Klosen et al., 2013) and appetite regulator pro-opiomelanocortin (POMC) (Reddy et al., 1999; Mercer et al., 2000, 2001; Rousseau et al., 2002; Bao et al., 2019). This neuroendocrine pathway is acting as a seasonal on/off switch of the gonadotropic and metabolic axes.

Although these seasonal responses are well conserved among individuals and species, they may be threatened by new environmental constraints emerging from modern lifestyles. Notably, ubiquitous exposure to chemicals with endocrine-disrupting activities, called endocrine-disrupting chemicals (EDCs), may impact the seasonal neuroendocrine mechanisms, an issue still not investigated in seasonal mammals (Moralia et al., 2022). Yet, bisphenol A (BPA), a well-described EDCs massively used by the plastic and resin industries, displays oestrogenic (Kuiper et al., 1998; Kitamura et al., 2005; Delfosse et al., 2012), anti-androgenic (Wang et al., 2017), and anti-thyroid hormone activities (Moriyama et al., 2002) which could alter the complex neuroendocrine control of seasonal physiology. Moreover, BPA has been reported to modify the activity of kisspeptin, RFRP-3 (López-Rodriguez et al., 2021), and POMC neurons (Desai et al., 2018; Salehi et al., 2019) that are involved in the seasonal synchronization of the gonadotropic and metabolic axes.

In this study, we tested the hypothesis that BPA exposure impacts the reproductive and metabolic adaptations to photoperiodic changes in Djungarian hamsters, with putative sex differences.

## 2. Material and methods

### 2.1. Animals

This project involved sexually mature (6 month-old on average) male and female Djungarian hamsters, born and raised in the Chronobiotron animal facility (UMS 3415, Strasbourg, France) at a temperature of 22 ± 1°C under LP (16 h light/8 h dark) conditions. Hamsters were grouped by 3 or 4 in type II L cages, enriched with nest cottons, wooden sticks, and a stainless steel tunnel, with *ad libitum* access to food and water. To avoid as much as possible environmental BPA contamination, care was taken to place animals in polyphenylsulfone cages equipped with glass drinking bottles. They were fed with low phytoestrogen diet (7011 P, Altromin International) prepared using glassware, stainless steel, and polypropylene tubes. All animal experiments were approved by the local ethical committee on animal experimentation (CREMEAS), in accordance with the French ministry of Higher Education and Scientific Research (authorization #22595-2019102317455140).

### 2.2. Experimental design

Animals were first adapted to eat the reconstituted low phytoestrogen diet for at least 6 weeks. From the start of both experimental protocols (experiment 1: transfer from LP to SP (8h light/16 h dark) or experiment 2: transfer from SP to LP), hamsters were fed with the reconstituted low phytoestrogen diet containing either 0.02% ethanol (control animals with BPA solvent) or BPA (#239658, Sigma Aldrich) adjusted to a dose of 5, 50, or 500 μg/kg of body weight/day (BPA/kg/day). For each group of animals, BPA quantity added to the reconstituted food was adjusted each week according to the average body weight and amount of food intake measured the week before.

**Experiment 1** assessed the effects of BPA exposure on the reproductive and metabolic integration of SP in male and female Djungarian hamsters. Hamsters raised in LP were transferred to SP for 10 weeks and fed with a diet containing different doses of BPA (0, 5, 50, or 500 μg of BPA/kg/day, n=5 to 8 per sex and experimental group, **figure 1A**). Another group of hamsters (n=8 to 10 per sex) was maintained in LP conditions without BPA to serve as a reference group for active reproduction and high metabolic activity. During the 10-week SP adaptation, hamster’s food intake, body weight, and fur color were monitored, and at the end of this experiment, they were sacrificed as explained below for further analyses.

**Figure 1.**
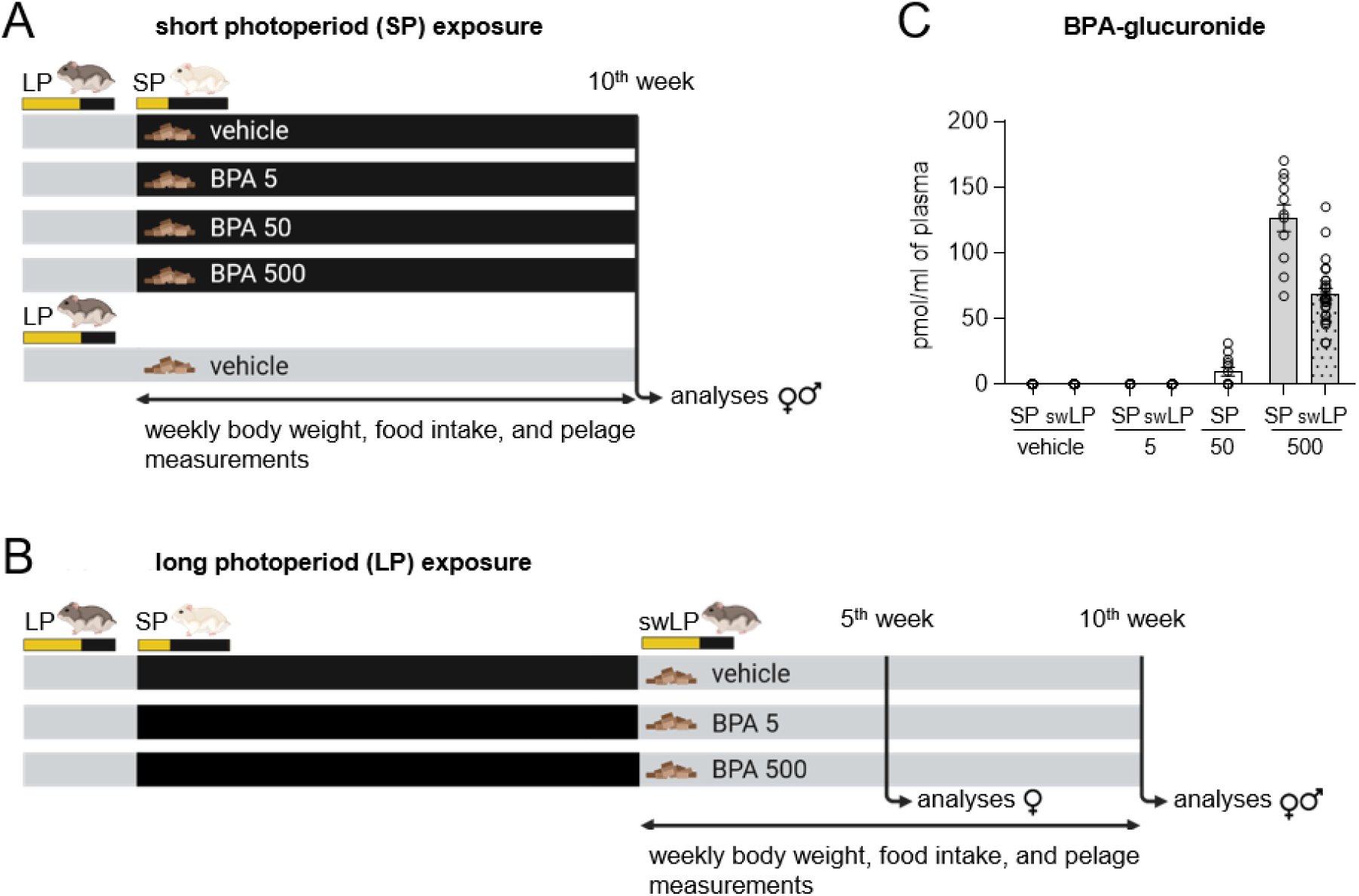
Paradigms of exposure to BPA during photoperiodic transfers and plasmatic BPA-glucuronide concentrations. Female and male hamsters were exposed to the vehicle (0.02% ethanol) or BPA at 5 (BPA 5), 50 (BPA 50), or 500 (BPA 500) µg/kg/day incorporated in food for 10 weeks of a long (LP) to short (SP) photoperiod transfer (**A**) or a SP to LP transfer (swLP) (**B**). Body weight, food intake, and pelage score were monitored each week, and various tissues were sampled at the end of the 10-week experiment (except for an additional group of females sampled 5 weeks after the SP to LP transfer). The BPA metabolite, BPA-glucuronide, was measured in the plasma of each animal to confirm the effectiveness of the administration route (**C**).

**Experiment 2** assessed the effects of BPA exposure on the male and female Djungarian hamster’s reproductive and metabolic reactivation induced by a switchback from SP to LP (called swLP). Hamsters were first adapted to SP for 11 weeks, and animals exhibiting the expected reproductive inactivation and body weight loss were transferred in LP conditions for 10 weeks and fed with a diet containing different doses of BPA (0, 5, or 500 μg of BPA/kg /day, n=7 to 12 per sex and experimental group, **figure 1B**). During the LP adaptation, hamster’s food intake, body weight, and fur color were monitored. Half females were sacrificed 5 weeks, and the second half 10 weeks after LP transfer. A considerable death of male hamsters due to increased aggressive behavior in SP (Jasnow et al., 2000) did not allow us to separate the males into two longitudinal sampling groups, and they were all sacrificed 10 weeks after LP transfer.

Of note and as already reported (Gorman and Zucker, 1997), some individuals maintained a LP-like reproductive and metabolic phenotype after the SP transfer, as they did not exhibit the expected body weight loss, gonadal atrophy, or fur whitening. In our experiment, and according to the exclusion criteria established in the literature (Butler et al., 2007; Greives et al., 2008), 5 hamsters out of 148 were identified as photoperiodic non-responders and were thus removed from all the analyses.

### 2.3. Body weight, food intake and fur color analyses

Individual hamster’s body weight was measured once a week. A non-linear regression model was used to determine the timing and speed of the photoperiodic changes in body weight, as described before (Butler et al., 2007). For each hamster, either a SP dose-response inhibition equation (experiment 1) or a LP dose-response activation equation (experiment 2) was fitted to the normalized body weight values (mean R² = 0.91 for experiment 1; mean R² = 0.85 for experiment 2) to determine acceleration in body weight change (i.e. the point of inflection of the body weight loss or body weight gain curves), and Hill slope. For experiment 1, transition to the SP body weight phenotype was defined as the point of zero acceleration in the period of body weight loss, i.e. the point of inflection in the body weight loss curve. For experiment 2, transition to the LP body weight phenotype was defined as the point of maximum acceleration in the period of body weight gain, i.e. the point of inflection in the body weight gain curve. Speeds of body weight loss or gain were estimated from Hill slopes.

Food intake was measured once a week per cage of 3 to 4 hamsters. Cumulative feeding efficiency ratios were determined by dividing cumulative individual body weight gain or loss by the average cumulative food intake of each cage, over all weeks of experimental treatment.

The change in fur coloration and density was scored weekly using an index described by Figala et al (1973). The score ranged from 1 for brown summer fur to 5 for white winter fur.

### 2.4. Tissue sampling

At the end of experiments, all hamsters were sacrificed during the early light phase (between 1 to 4 hours after lights on) with the females being sacrificed in diestrus to reduce hormonal bias. Each hamster was deeply anesthetized (40 mg/kg of tiletamine/zolazepam (Zoletil^®^50) and 10 mg/kg of xylazine (Paxman)), then blood was collected by heart puncture, and the body was fixed by a transcardiac perfusion with phosphate buffered saline (PBS) followed by 4% paraformaldehyde-lysine-periodate (PLP) fixative. After perfusion, reproductive organs (uterus with ovaries for females and pairs of testes for males) were directly dissected and weighed. Brains were extracted and conserved in the PLP solution for 12 hours before dehydration and polyethylene glycol embedding (Klosen et al., 1993). Each brain was cut into 10 µm-thin coronal sections from the preoptic area to the mammillary bodies, and 10 sets of 16 serial sections covering the full extent of this hypothalamic area were mounted on Superfrost Plus glass slides, selecting one out of every 10 sections (consecutive sections being 100 µm apart). Thus, each glass slide contained 16 sections spanning the hypothalamus of one animal. Slides were dried 20 minutes at 50°C and stored at –80°C until use for neuroanatomical analyses.

### 2.5. Measurements of circulating BPA metabolite, sex steroids, glucose, and metabolic hormones

**Plasma testosterone and BPA-glucuronide** were analyzed by LC/MS-MS. Briefly, 15 µL of internal standards (3.3 µM –[^2^H4]-testosterone + 13.3 µM [^13^C12]-BPA-glucuronide) and 450 µL of acetonitrile (ACN) were mixed to 150 µL of plasma sample. After centrifugation at 20,000xg for 30 minutes at 4°C, the supernatant was collected, dried under vacuum, and resuspended in 30 µL of 40% ACN/0.1% formic acid (v/v), and centrifuged at 20,000xg for 20 minutes at 4°C. The final supernatant was divided in two samples of 15 µL for testosterone and 15 µL for BPA-glucuronide analyses. For testosterone assay only, the 15 µL supernatant was dried under vacuum and resuspended in 15 µL of Amplifex Keto Reagent Kit solution (Waters) for 1 hour at room temperature to enhance testosterone LC/MS-MS signal, and finally centrifuged at 20,000xg for 10 minutes at 4°C. Analyses were conducted on 4 µl of both samples with a Dionex Ultimate 3000 HPLC system coupled with a triple quadrupole Endura mass spectrometer (Thermo Electron) in the positive (testosterone) or negative mode (BPA-glucuronide). Separation of the testosterone and BPA-glucuronide were done at 40°C on a Zorbax column with the gradient of ACN. The identification of the compounds was based on precursor ions, selective fragment ions (i.e. daughter ions) and retention times obtained for testosterone, BPA-glucuronide, and their internal standards. Quantification was done by the calculation of the peak area ratio between the daughter ion used for quantification of testosterone or BPA-glucuronide and their internal standards (technical information are in supplementary Tables 1 and 2).

The measurement of plasma BPA-glucuronide, the major metabolite of BPA, allowed to confirm that BPA exposure induced relevant concentrations of BPA in the organism of exposed animals, and to verify that control animals were exposed to minimal levels of environmental BPA (Figure 1C). The absence of BPA-glucuronide detection in the plasma of the 5 µg BPA/kg/day-exposed animals may be due to a limit of sensitivity of our assay.

**Plasma estradiol** was measured after extraction in methanol according to the Estradiol ELISA Kit (#501890, Cayman Chemical). Intra– and inter-assay variations were < 5%, and the assay displayed a sensitivity of 20 pg/mL.

**Plasma insulin** was measured in 20 times diluted plasma samples using the Hamster Insulin ELISA Kit (#90336, CrystalChem). Intra-assay variations were < 10% and inter-assay variations were < 15%. The assay has a limit of sensitivity of 0.05 ng/mL.

**Blood glucose** levels were measured immediately in one drop of blood after the heart puncture with the Blood Glucose System Accu-Chek Performa (Roche Diagnostics).

### 2.6. Neuroanatomical analyses

For each staining, all slides of all animals belonging to the same experimental protocol were processed together. **Figure 2** illustrates the neuroanatomical localisation of the investigated genes and protein.

**Figure 2.**
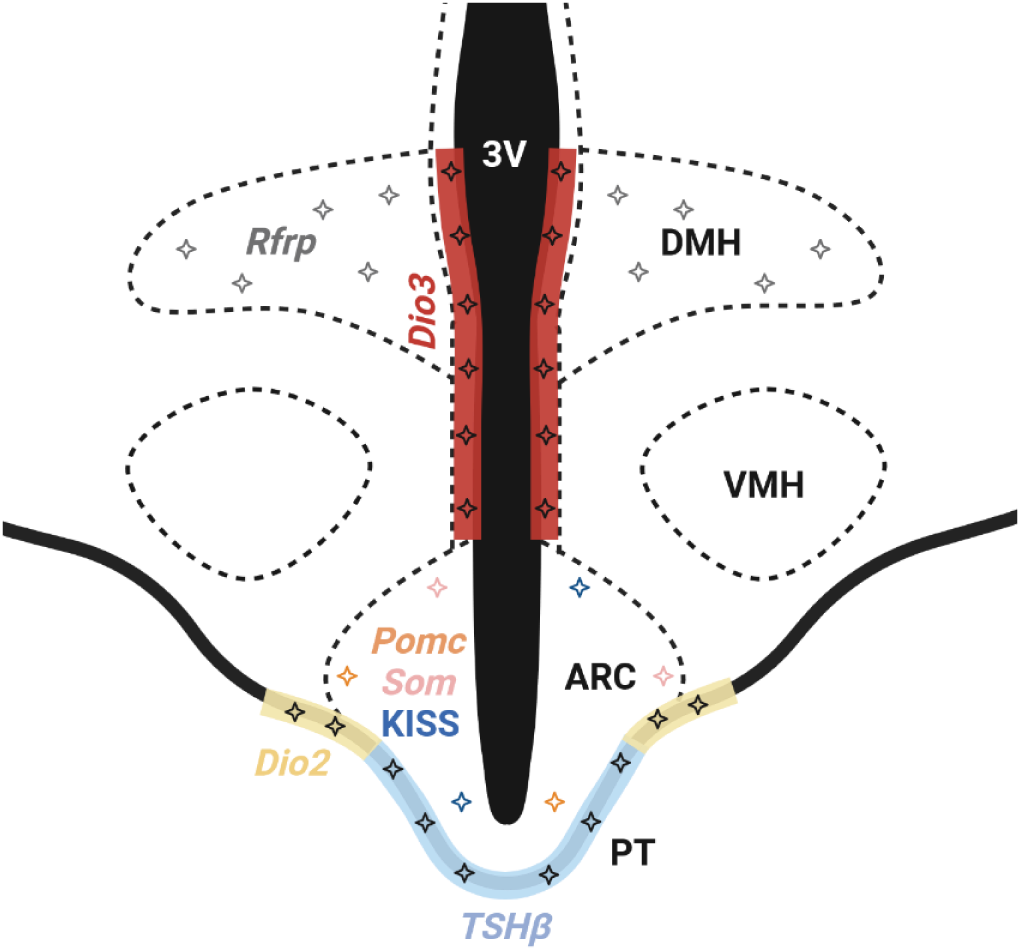
Representation of the anatomical location of the photoperiodic genes and proteins analyzed using histological techniques. The mRNA or protein expression of *Pomc*, *somatostatin* (*Som*), and kisspeptin (KISS) was quantified in the arcuate nucleus (ARC), *Rfrp* in the dorsomedial hypothalamus (DMH). *Dio2* analysis was performed on the tanycyte endfeets in the tuberoinfundibular sulcus, while *Dio3* was analyzed alongside the third ventricle (3V) in the α-tanycytes body cells. *TSHβ* expression was analyzed alongside the *pars tuberalis* (PT). VMH = ventromedial hypothalamus.

**Non-radioactive *in situ* hybridization** was performed on fixed hypothalamic sections using antisens riboprobes transcribed from 1 µg linearized plasmid DNA with digoxigenin or fluorescein-labelled nucleotides. Djungarian hamster’s *Dio2*, *Dio3* and *Rfrp* probes were used, as well as rat *TSHβ*, *Pomc*, and *somatostatin* probes that were all previously validated on Djungarian hamster tissue (Milesi et al., 2017; Cázarez-Márquez et al., 2019).

Briefly, the sections were post-fixed for 10 minutes with 4% formalin in phosphate buffer (PB), rinsed in PBS, digested for 30 minutes at 37°C with 0.5-1 µg/ml of proteinase K (Roche), quickly rinsed with cold PBS, post-fixed with 2% formalin in PB on ice, rinsed in PBS + 0.01% diethylpyrocarbonate (DEPC), acetylated twice for 10 minutes in 100 mM triethanolamine + 0.25% acetic anhydride, washed in PBS + 0.01% DEPC and equilibrated in 5X saline sodium citrate (SSC) + 0.05% Tween 20 + 0.01% DEPC. The sections were hybridized with 200 ng/ml denaturated riboprobes diluted in 50% formamide + 5X SSC + 5X Denhardt’s solution + 1 mg/ml of salmon DNA + 0.1% Tween 20 + 0.04% DEPC for 40 hours at 54-60°C. After hybridization, sections were washed in SSC 5X + 0.05% Tween 20 and then in SSC 0.1X + 0.05% Tween 20 at 72°C to remove non-specific hybrids. Sections were washed in A-DIG buffer + 0.05% Tween 20, blocked for at least 1 hour with blocking buffer (Roche), and incubated overnight with the alkaline phosphatase (AP) coupled anti-digoxigenin or anti-fluorescein antibody (Roche) at respectively 1/5000 or 1/1000 in blocking buffer. Sections were rinsed in A-DIG buffer + 0.05% Tween 20 and equilibrated in AP 1X buffer. AP activity was detected either with a solution of nitro blue tetrazolium (Roche) / bromo-chloro-indolyl phosphate (Thermo Scientific) in AP buffer or using Naphtol AS-MX phosphate and Fast Red (Sigma-Aldrich) as substrate for 2 to 24 hours depending on the probes. Reactions were stopped with tap water before the staining intensity reached saturation. Slides were premounted with CrystalMount (Sigma-Aldrich), dipped in toluene, and mounted with a coverslip and Eukitt (Sigma-Aldrich).

**Immunohistochemistry**. Because *Kiss1* mRNA expression in the ARC is difficult to quantify in Djungarian hamsters, we used immunolabelling to detect ARC kisspeptin (Milesi et al., 2017). Brain sections were treated for antigen reactivation with citrate buffer at 95°C for 2 hours, cooled down to room temperature and rinsed with tris-buffered saline (TBS). Slides were pretreated with a blocking reagent (TBS + 3% powder milk + 0.02% NaN_3_) for at least 1 hour before being incubated overnight with the primary rabbit kisspeptin antibody (JLV-1 antiserum against the rat kisspeptin-52, Mikkelsen and Simonneaux, 2009) diluted at 1/1500 in TBS + 0.05% Tween 20 + 1% donkey serum + 0.02% NaN_3_. Sections were rinsed in TBS + 0.02% Tween 20, incubated for 1 hour with a biotinylated secondary antibody (donkey anti-rabbit (Jackson) diluted at 1/2000 in TBS + 0.02% Tween 20 + 1% donkey serum), rinsed in TBS + 0.02% Tween 20, and incubated with a neutravidin-horseradish peroxidase (HRP, Thermo Scientific) solution at 1/2000 in TBS + 0.02% Tween 20 + 0.2% cold water fish gelatin for 1 hour. Sections were washed in TBS + 0.02% Tween 20, equilibrated in tris-imidazole buffer (TBI), and HRP activity was detected with a solution of 1% diaminobenzidine (Acros Organics) + 0.003% H_2_O_2_ in TBI. Sections were dehydrated in alcohol baths and coverslipped with Eukitt.

**Image analysis**. For *somatostatin*, *Pomc*, *TSHβ*, and *Dio3* staining obtained from experiment 1, micrographs were taken at 10 × 0.63 objective by a DP 50 digital camera (Olympus) attached to a DMRB microscope (Leica). A background image without tissue was taken for each slide and subtracted from sampled images. The images were analyzed using ImageJ software (Rasband, U.S. National Institute of Health). Mean gray value pixels of *somatostatin* and *Pomc* labelling in neurons were measured using a fixed-size circle overlaid on at least 50 neurons per animal, as described before (Klosen et al., 2013; Cázarez-Márquez et al., 2019). As *somatostatin* mRNA expression exhibits photoperiodic variations only in the caudal part of the ARC (Herwig et al., 2012), *somatostatin* neuron labelling intensity was measured only in 3-4 images/animal of this region. *Pomc* neuron labelling intensity was measured in 6-7 images/animal covering the full extent of the ARC. *TSHβ* integrated density was measured in 3-4 images/animal using a 65-µm wide line tool covering the *pars tuberalis*. For *Dio3*, we used standardized thresholding to select and measure integrated density using a 150-µm wide line tool in Dio3-stained areas (in α tanycytes) of 3 thresholded images/animal. *Rfrp* labelled neurons were observed with an optic microscope equipped with a camera, counted manually, and normalized by the number of sections counted. When mouting, we included sections that did not contained *Rfrp* labelling rostrally and caudally to ensure a total count of the number of hypothalamic *Rfrp* expressing neurons.

For kisspeptin staining and all stainings obtained from the experiment 2, slides were scanned with the Nanozoomer 2.0 HT (Hamamatsu) using the program NDP.scan. The imageJ analyses for *Dio2*, *TSHβ*, and *somatostatin* stainings was the same as described for the experiment 1. In order to improve consistency of counts and to detect all labelled cells when the number of cells is important, as it is for kisspeptin, *Pomc* and *Rfrp* expressing neurons, an automatic cell detection was performed using QuPath software (v0.4.0, Bankhead et al., 2017). We used the StarDist deep-learning-based method of cell detection to count kisspeptin labeled cells, based on the pre-trained model ‘dsb2018_heavy_augment.pb’ (Schmidt et al., 2018). *Pomc* and *Rfrp* labelled neurons were detected automatically based on morphological and intensity criteria, with post-checking that the detected objects were indeed labelled cells of interest.

### 2.7. Statistical analysis

All data are presented as mean ± SEM of n animals (from 5 to 12 according to the experimental groups) and were analyzed using GraphPad Prism 8.0.2. Repeated values over time were analyzed by Two-Way analysis of variance (ANOVA) followed by Dunnett’s *post hoc* tests. Mixed-model effects were used instead of Two-Way ANOVA when values were missing. For the females of the experiment 2, for each dose, the dynamic changes of mRNA/protein expression between the 5^th^ and the 10^th^ week were analyzed using Two-Way ANOVA with Sidak *post hoc* tests for multiple comparisons. Multiple group comparisons were analyzed using One-Way ANOVA or Kruskal-Wallis tests, followed by Dunnett’s *post hoc* test. To ascertain the photoperiodic effect, comparisons between the LP– and SP– or swLP-vehicle groups were performed using t-tests. Statistical significance was set at p-value < 0.05.

## 3. Results

### 3.1. Effect of oral BPA exposure on the physiological and neuroendocrine short photoperiod integration in female and male hamsters

#### 3.1.1. Exposure to a high dose of BPA delays metabolic integration in female hamsters under short photoperiod

All female hamsters showed an expected decrease in body weight upon transfer to SP, while those kept in LP maintained stable body weight (**Figure 3A**). However, females exposed to 500 µg BPA/kg/day displayed a delayed SP-induced body weight loss by more than one week compared to those exposed to the vehicle, 5, or 50 µg of BPA/kg/day (**Figure 3B**). Furthermore, females exposed to 500 µg BPA/kg/day demonstrated higher feed efficiency ratios (FER) during the initial two weeks of SP compared to the vehicle-exposed group (**Figure 3C**). After 10-week SP exposure, all female hamsters lost an average of 24.6% (± 1.1%) of their body weight whether exposed or not to BPA (**Figure 3B**). The glucose/insulin (G/I) ratio (**Table 1**), an indicator of insulin sensitivity, was similar among all female groups. Fur color whitening was also delayed in females exposed to 500 µg BPA/kg/day as compared to the vehicle-exposed group, although all females exhibited a similar white fur color 10 weeks after the SP transfer **(Figure 3D**).

**Figure 3.**
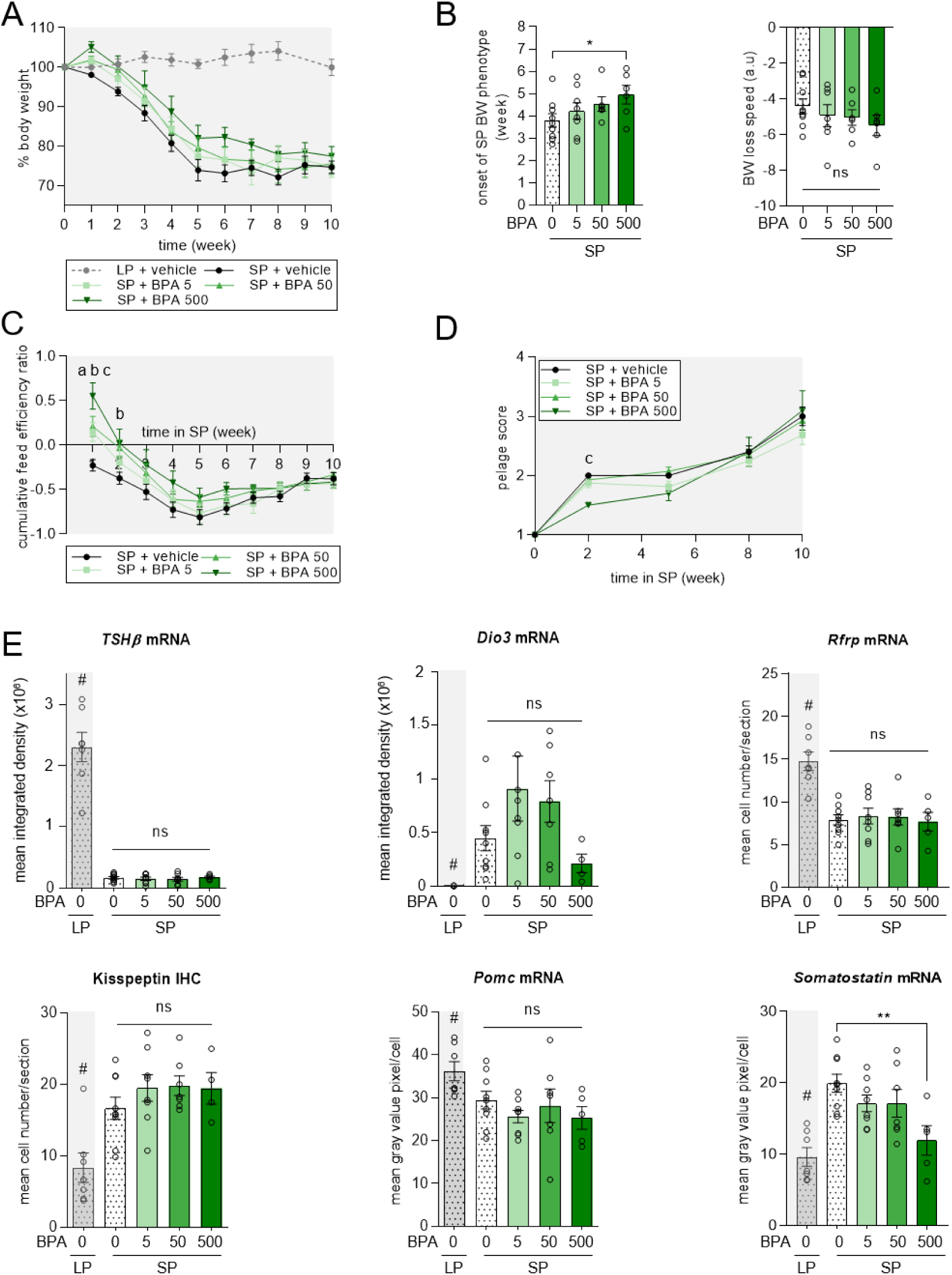
Effects of BPA exposure on body weight, food intake, pelage whitening, and various photoperiodic genes/proteins in female Djungarian hamsters transferred from long to short photoperiod. Female Djungarian hamsters were either kept in long photoperiod (LP) and fed with pellet food containing 0.02% ethanol (LP + vehicle; n=8) or transferred in short photoperiod (SP) for 10 weeks and fed with pellet food containing 0.02% ethanol (SP + vehicle; n=9), or containing 5 (SP + BPA 5; n=8), 50 (SP + BPA 50; n=7), or 500 (SP + BPA 500; n=6) µg BPA/kg/day. **(A)** Body weight (BW) expressed in percentage of the LP value (before the SP transfer) during the 10 weeks of photoperiodic treatment. **(B)** Time in weeks to reach the SP body weight phenotype (point of inflection calculated from a non-linear regression model fitted to individual body weight raw data) and speed of SP-induced body weight loss (slope of the non-linear regression). **(C)** Cumulative feed efficiency ratio over 10 weeks of SP. **(D)** Pelage score (from 1 being a LP-adapted grey color to 5 being a SP-adapted white color) over 10 weeks of SP. **(E)** Expression of *pars tuberalis TSHβ*, and hypothalamic *Dio3*, *Rfrp*, *Pomc*, and *somatostatin* gene and kisspeptin protein in constant LP or after 10 weeks of SP. Values are given as mean ± SEM (n=6 to 9 according to experimental groups). Statistical significance: for repeated values over time, *a*, p<0.05 between SP + BPA 5 vs SP + vehicle, *b*, p<0.05 between SP + BPA 50 vs SP + vehicle, *c*, p<0.05 between SP + BPA 500 vs SP + vehicle; for multiple comparisons between groups, *ns*, no significance; *, p<0.05 vs SP + vehicle; **, p<0.01 vs SP + vehicle; to ascertain the photoperiodic effect in gene/protein expression, #, p<0.05 between LP + vehicle vs SP + vehicle.

**Table 1.** Effects of BPA exposure on reproductive and metabolic parameters in female and male Djungarian hamsters transferred from long to short photoperiod. Hamsters were orally exposed (through food) to 0.02% ethanol (SP + vehicle; n=8-9) or to 5 (SP + BPA 5; n=7-8), 50 (SP + BPA 50; n=6-7), or 500 (SP + BPA 500; n=5-6) µg of BPA/kg/day during a 10-week transfer from a long (LP) to a short photoperiod (SP). A control group of hamster was kept under LP (LP + vehicle; n=8-10). Values are indicated as mean ± SEM. Statistical significance: #, p<0.05 between LP + vehicle vs SP + vehicle, *a*, p<0.05 between SP + BPA 5 vs SP + vehicle. ND = non detectable

|  | females |  |  |  |  | males |  |  |  |  |
| --- | --- | --- | --- | --- | --- | --- | --- | --- | --- | --- |
| photoperiod/dose | LP<br>+ vehicle | SP<br>+ vehicle | SP<br>+ BPA 5 | SP<br>+ BPA 50 | SP<br>+ BPA 500 | LP<br>+ vehicle | SP<br>+ vehicle | SP<br>+ BPA 5 | SP<br>+ BPA 50 | SP<br>+ BPA 500 |
| **relative reproductive organ weight**<br>(mg/g of body weight) | 2.51 ± 0.21<sup>#</sup> | 1.74 ± 0.06 | 1.96 ± 0.15 | 1.78 ± 0.15 | 1.93 ± 0.22 | 20.30 ± 1.76<sup>#</sup> | 1.77 ± 0.25 | 1.60 ± 0.14 | 1.56 ± 0.18 | 1.50 ± 0.14 |
| **estradiol**<br>(pg/ml of plasma) | 283.4 ± 63.2 | 232.0 ± 114.7 | 337.1 ± 125.1 | 162.4 ± 45.7 | 146.8 ± 44.5 |  |  |  |  |  |
| **testosterone**<br>(pg/ml of plasma) |  |  |  |  |  | 35.81 ± 15.70 | ND | ND | ND | ND |
| **glucose/insulin ratio** | 19.40 ± 4.13 | 24.43 ± 2.88 | 27.17 ± 4.19 | 25.69 ± 5.27 | 24.8 ± 3.44 | 14.42 ± 3.78 | 23.54 ± 5.45 | 49.15 ± 10.16<sup>a</sup> | 40.12 ± 11.34 | 34.36 ± 5.15 |

At the end of the 10-week SP exposure, all female hamsters exhibited the expected decrease in the relative uterine and ovary weight, with no significant effect of BPA exposure on the final regression of the reproductive organs (**Table 1**). Circulating concentrations of estradiol were similar among all female hamster groups.

The selected genes or proteins known to adapt reproductive and metabolic activities to photoperiodic changes showed the expected SP-induced decrease (*TSHβ*, *Rfrp*) and increase (*Dio3,* kisspeptin, *somatostatin*) in their expression 10 weeks after the SP transfer, compared to LP hamsters (**Figure 3E**). However, exposure to 500 µg BPA/kg/day abolished the SP-induced increase in ARC somatostatin. The photoperiodic changes in the other investigated genes or protein were not altered by BPA exposure.

#### 3.1.2. Exposure to a low dose of BPA accelerates metabolic integration in male hamsters under short photoperiod

All male hamsters switched to SP gradually reduced their body weight (**Figure 4A**). However, the use of a non-linear regression model indicates that the onset of the SP body weight phenotype appeared one week earlier in hamsters exposed to 5 µg BPA/kg/day compared to hamsters exposed to vehicle, 50, or 500 µg BPA/kg/day (**Figure 4B**). The feed efficiency ratio was also lower in male hamsters exposed to 5 µg of BPA/kg/day compared to the vehicle-exposed group 5 weeks after the SP transfer (**Figure 4C**), and the G/I ratio was higher for the 5 µg BPA/kg/day-exposed group compared to the other experimental groups (**Table 1**). The speed of the SP-induced body weight loss was similar among all groups of male hamsters, and all lost an average of 23.9% (± 1.4%) of their body weight 10 weeks after the SP transfer, whether exposed or not to BPA (**Figure 4B**).The pelage bleaching was significantly advanced in males exposed to 5 and 50 µg BPA/kg/day at the 8th week of SP, but after 10 weeks of SP exposure, all hamsters exhibited the same white and fluffy fur, whether exposed or not to BPA (**Figure 4D**).

**Figure 4.**
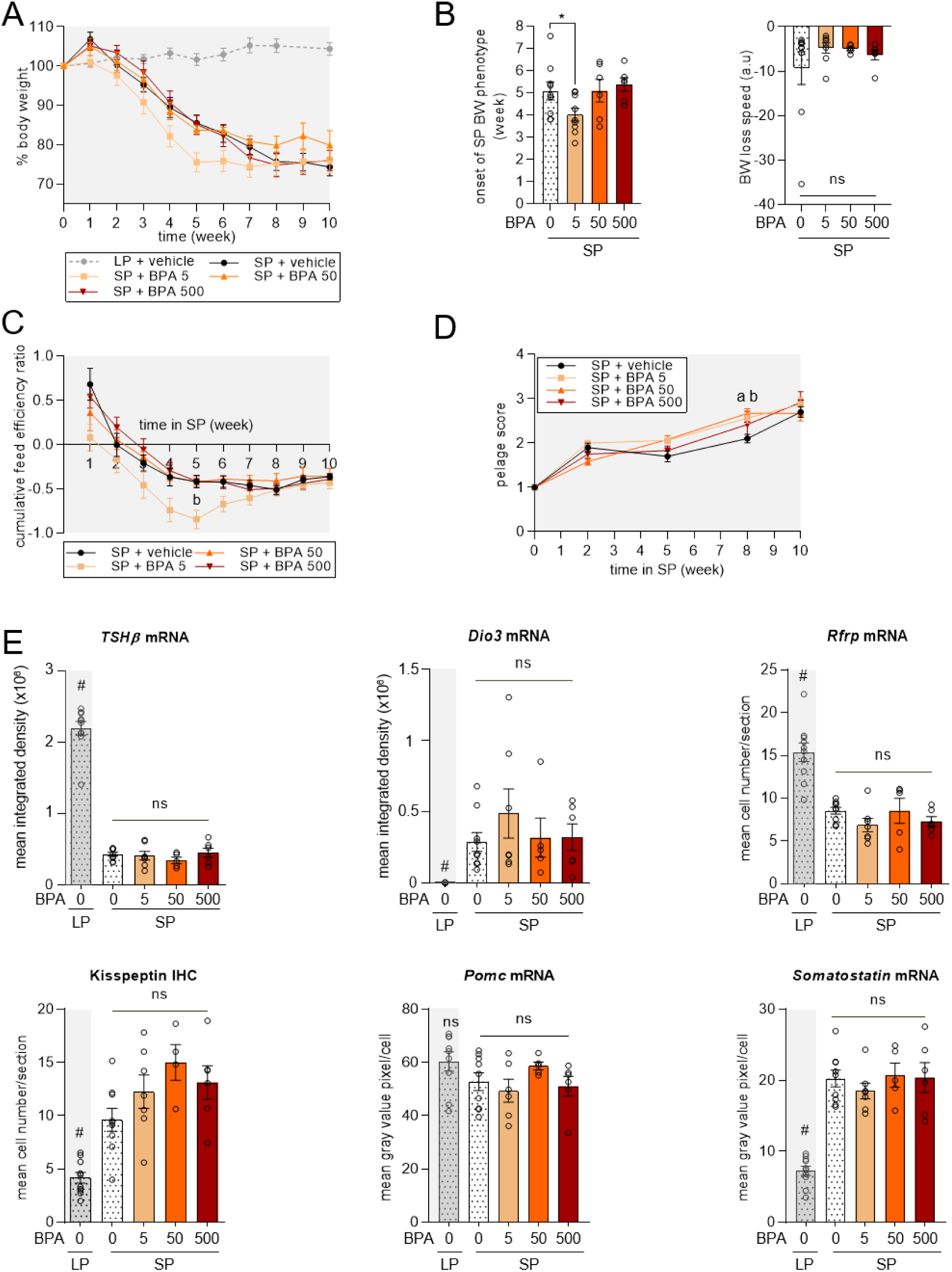
Effects of BPA exposure on body weight, food intake, pelage whitening, and various photoperiodic genes/proteins in male Djungarian hamsters transferred from long to short photoperiod. Male Djungarian hamsters were either kept in long photoperiod (LP) and fed with pellet food containing 0.02% ethanol (LP + vehicle; n=10), or transferred in short-photoperiod (SP) and fed with a diet containing 0.02% ethanol (SP + vehicle; n=9), or containing 5 (SP + BPA 5; n=8), 50 (SP + BPA 50; n=6), or 500 µg BPA/kg/day (SP + BPA 500; n=6). **(A)** Body weight (BW) expressed in percentage of the LP value (before the SP transfer) during the 10 weeks of photoperiodic treatment. **(B)** Time in weeks to reach the SP body weight phenotype (point of inflection calculated from a non-linear regression model fitted to individual body weight raw data) and speed of SP-induced body weight loss (slope of the non-linear regression).**(C)** Cumulative feed efficiency ratio over 10 weeks of SP **(D)** Pelage score (from 1 being a LP-adapted grey color to 5 being a SP-adapted white color) over 10 weeks of SP. **(E)** Expression of *pars tuberalis TSHβ*, and hypothalamic *Dio3*, *Rfrp*, *Pomc*, and *somatostatin* gene and kisspeptin protein after 10 weeks of SP or constant LP. Values are given as mean ± SEM (n=6 to 10 according to experimental groups). Statistical significance: for repeated values over time, *a*, p<0.05 between SP + BPA 5 vs SP + vehicle, *b*, p<0.05 between SP + BPA 50 vs SP + vehicle; for multiple comparisons between groups, *ns*, no significance; *, p<0.05 vs SP + vehicle; to ascertain the photoperiodic effect in gene/protein expression, #, p<0.05 between LP + vehicle vs SP + vehicle.

After 10 weeks of SP, all male hamsters had low testis weight, compared to hamsters kept in LP, and BPA exposure did not affect the final SP-induced regression of reproductive organs (**Table 1**). Plasma testosterone was only detectable in male hamsters kept in LP and was below the limit of detection for all SP-adapted groups.

As for female hamsters, all of the selected reproductive and metabolic genes or proteins showed the expected SP-induced changes, and exposure to various doses of BPA did not modify the final SP-induced changes in these gene and protein expressions (**Figure 4E**).

### 3.2. Effect of oral BPA exposure on the physiological and neuroendocrine integration of long photoperiod in male and female hamsters

After a prior full adaptation to 11 weeks of SP exposure, male and female hamsters were switched back to LP (swLP) with or without oral exposure to BPA for a subsequent 10-week period. The 50 µg/kg/day BPA dose was excluded in this second study based on observations from the previous experiment, where the 5 and 500 µg/kg/day doses showed more pronounced physiological and neuroendocrine changes. Further, as the previous BPA exposure was found to alter the kinetics of some aspects of photoperiodic adaptation, intermediate groups of hamsters were added to be sampled 5 weeks after the swLP transfer. However, due to a large number of male hamster deaths, only female hamsters were available for sampling at this intermediate stage.

#### 3.2.1. BPA exposure induces a faster metabolic and reproductive adaptation to long photoperiod in female hamsters

All females transferred back to LP conditions gradually regained body weight (**Figure 5A**), although females exposed to 5 and 500 µg BPA/kg/day experienced a faster body weight gain (**Figure 5B**) and a higher increase in feed efficiency ratios in the initial two weeks of swLP (**Figure 5C**) compared to vehicle-exposed females. After 10 weeks in swLP, all females gained an average of +23.1% (± 4.7%) body weight, regardless of BPA exposure (**Figure 5A**), and blood glucose concentrations were comparable across all experimental groups (**Table 2**). Throughout the 10 weeks of swLP, female hamster coats remained whitish, with no observable differences among groups (**Figure 5D**).

**Figure 5.**
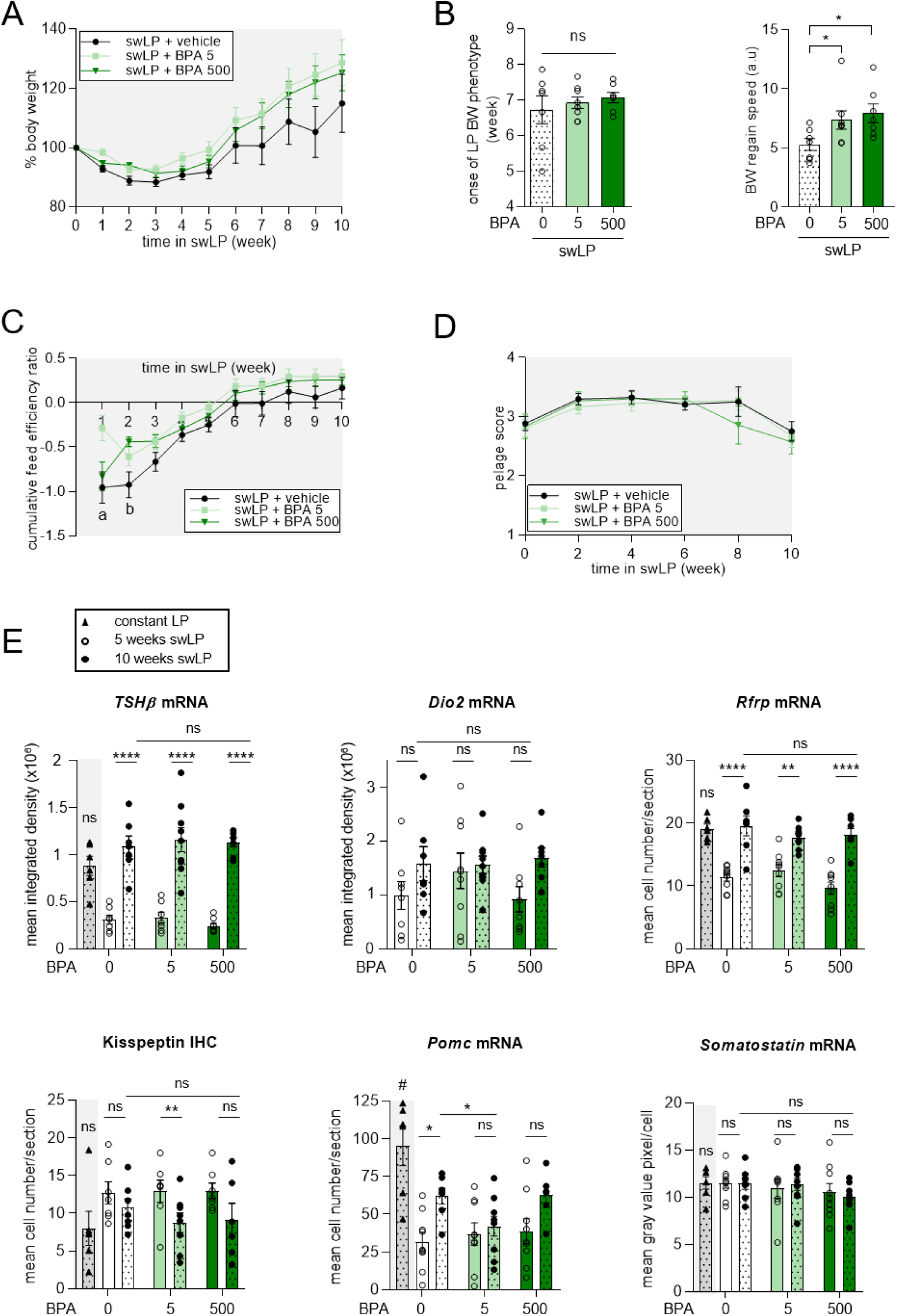
Effects of BPA exposure on body weight, food intake, pelage whitening, and various photoperiodic genes/proteins in female Djungarian hamsters transferred from short to long photoperiod. Female Djungarian hamsters adapted to short photoperiod (SP) were switched back in long photoperiod (swLP) and either fed with pellet food containing 0.02% ethanol (swLP + vehicle; n=8-17), or containing 5 (swLP + BPA 5; n=9-18) or 500 µg (swLP + BPA 500; n=7-16) BPA/kg/day. **(A)** Body weight (BW) expressed in percentage of the SP value (before the swLP transfer). **(B)** Time in weeks to reach the LP body weight phenotype (point of inflection calculated from a non-linear regression model fitted to individual body weight raw data) and speed of LP-induced body weight gain (slope of the non-linear regression). **(C)** Cumulative feed efficiency ratio over 10 weeks of swLP. **(D)** Pelage score (from 1 being a LP-adapted grey color to 5 being a SP-adapted white color) over 10 weeks of swLP. **(E)** Expression of *pars tuberalis TSHβ*, and hypothalamic *Dio2*, *Rfrp*, *Pomc*, and *somatostatin* gene and kisspeptin protein in constant LP or after 5 and 10 weeks of swLP., Values are given as mean ± SEM (n=7 to 18 according to experimental groups). Statistical significance: for repeated values over time, *a*, p<0.05 between swLP + BPA 5 vs swLP + vehicle, *b*, p<0.05 between swLP + BPA 50 vs swLP + vehicle; for multiple comparisons between groups at the 10^th^ week (B), *, p<0.05 vs swLP + vehicle; for multiple comparisons of the dynamic changes of mRNA/protein expression between the 5^th^ and the 10^th^ week for each dose, *, p<0.05, **, p<0.01, ***, p<0.001, ****, p<0.0001; to ascertain the photoperiodic effect in gene/protein expression, #, p<0.05 between LP + vehicle and swLP + vehicle. *ns*, no significance.

**Table 2.** Effects of BPA exposure on reproductive and metabolic parameters in female and male Djungarian hamsters transferred from short to long photoperiod. Hamsters were orally exposed (through food) to 0.02% ethanol (swLP + vehicle; n=8-11), or to 5 (swLP + BPA 5; n=9-11) or 500 (swLP + BPA 500; n=7-12) µg of BPA/kg/day after a 10-week transfer from a short to a long photoperiod (swLP), or when kept under long photoperiod conditions (LP + vehicle; n=8-10). Values are indicated as mean ± SEM. Statistical significance: #, p<0.05 between LP + vehicle vs swLP + vehicle (10^th^ week), *a*, p<0.05 between swLP + BPA 5 vs swLP + vehicle.

|  | females |  |  |  |  |  |  |  | males |  |  |
| --- | --- | --- | --- | --- | --- | --- | --- | --- | --- | --- | --- |
| photoperiod/<br>dose | LP + vehicle | swLP + vehicle |  | swLP<br>+ BPA 5 |  | swLP<br>+ BPA 500 |  | LP + vehicle | swLP + vehicle | swLP<br>+ BPA 5 | swLP<br>+ BPA 500 |
| weeks | - | 5<sup>th</sup> | 10<sup>th</sup> | 5<sup>th</sup> | 10<sup>th</sup> | 5<sup>th</sup> | 10<sup>th</sup> | - | 10<sup>th</sup> | 10<sup>th</sup> | 10<sup>th</sup> |
| relative<br>reproductive<br>organ weight<br>(mg/g of body) | 2.51<br>±<br>0.21<sup>#</sup> | 1.62<br>±<br>0.10 | 4.55<br>±<br>0.52 | 2.10<br>±<br>0.22<sup>a</sup> | 3.63<br>±<br>0.52 | 1.34<br>±<br>0.12 | 3.71<br>±<br>0.67 | 20.30<br>± 1.76 | 18.90<br>± 1.13 | 20.94<br>± 1.75 | 19.40<br>± 1.17 |
| estradiol<br>(pg/ml of<br>plasma) | 283.4<br>±<br>63.2 | 240.2<br>±<br>42.4 | 367.0<br>±<br>105.5 | 171.4<br>±<br>42.4 | 388.3<br>±<br>89.8 | 159.3<br>±<br>35.1 | 374.9<br>±<br>59.8 |  |  |  |  |
| testosterone<br>(pg/ml of<br>plasma) |  |  |  |  |  |  |  | 35.81<br>±<br>15.70 | 11.82<br>± 3.53 | 33.38<br>±<br>13.91 | 13.83<br>± 6.50 |
| glucose<br>(mg/dl of<br>blood) | 192.5<br>±<br>11.9 | 174.2<br>±<br>48.4 | 178.0<br>±<br>13.1 | 178.9<br>±<br>15.0 | 186.2<br>±<br>20.2 | 165.9<br>±<br>13.2 | 186.0<br>±<br>20.1 | 157.2<br>0 ±<br>6.16 | 156.0<br>0 ±<br>14.91 | 190.2<br>0 ±<br>18.72 | 146.2<br>0 ±<br>9.15 |

Five weeks after swLP, BPA-exposed females displayed significantly different relative uterine and ovarian weight compared to vehicle-exposed females (**Table 2**). Those exposed to 5 µg BPA/kg/day tended to have higher reproductive organ weight, while those exposed to 500 µg BPA/kg/day tended to have lower weight (One-way ANOVA, p<0.05, but no significant *post hoc* tests). After 10 weeks of swLP, all female groups reached a fully active reproductive state, as indicated by high reproductive organ weight and circulating estradiol levels (**Table 2**).

The comparison of mRNA/protein levels of most photoperiodic, metabolic, and reproductive genes and protein between the mid (5 weeks) and end (10 weeks) of swLP revealed the dynamic change during swLP integration (**Figure 5E**). Levels of the majority of investigated proteins/genes in 10-week swLP vehicle-treated females were similar to those of females maintained constantly in LP. However, exposure to 5 µg BPA/kg/day was associated with a significant decrease in kisspeptin level between the mid and end of swLP, suggesting a faster kisspeptin downregulation induced by the low BPA dose. Furthermore, in contrast to vehicle-treated hamsters, exposure to both doses of BPA abolished *Pomc* mRNA increase between the 5^th^ and the 10^th^ week of swLP, and exposure to 5 µg BPA/kg/day induced a lower *Pomc* mRNA expression at the 10^th^ week of swLP, suggesting that BPA delayed the LP-induced *Pomc* upregulation.

#### 3.2.2. BPA exposure during the long photoperiod adaptation of male hamsters alters the expression of photoperiodic gene but does not induce metabolic and reproductive effects

All male hamsters transferred from an SP to an LP condition gradually regained body weight, (approximatively +18.1 ± 3.4 % of initial (SP) body weight; **Figure 6A**), and increased their feed efficiency ratios (**Figure 6B**) without any significant impact of BPA exposure. Similarly, all hamster groups exhibited comparable blood glucose concentrations, independently of BPA exposure **(Table 2**). Throughout the 10 weeks of swLP, male hamster coats remained whitish, regardless of BPA exposure **(Figure 6C**).

**Figure 6.**
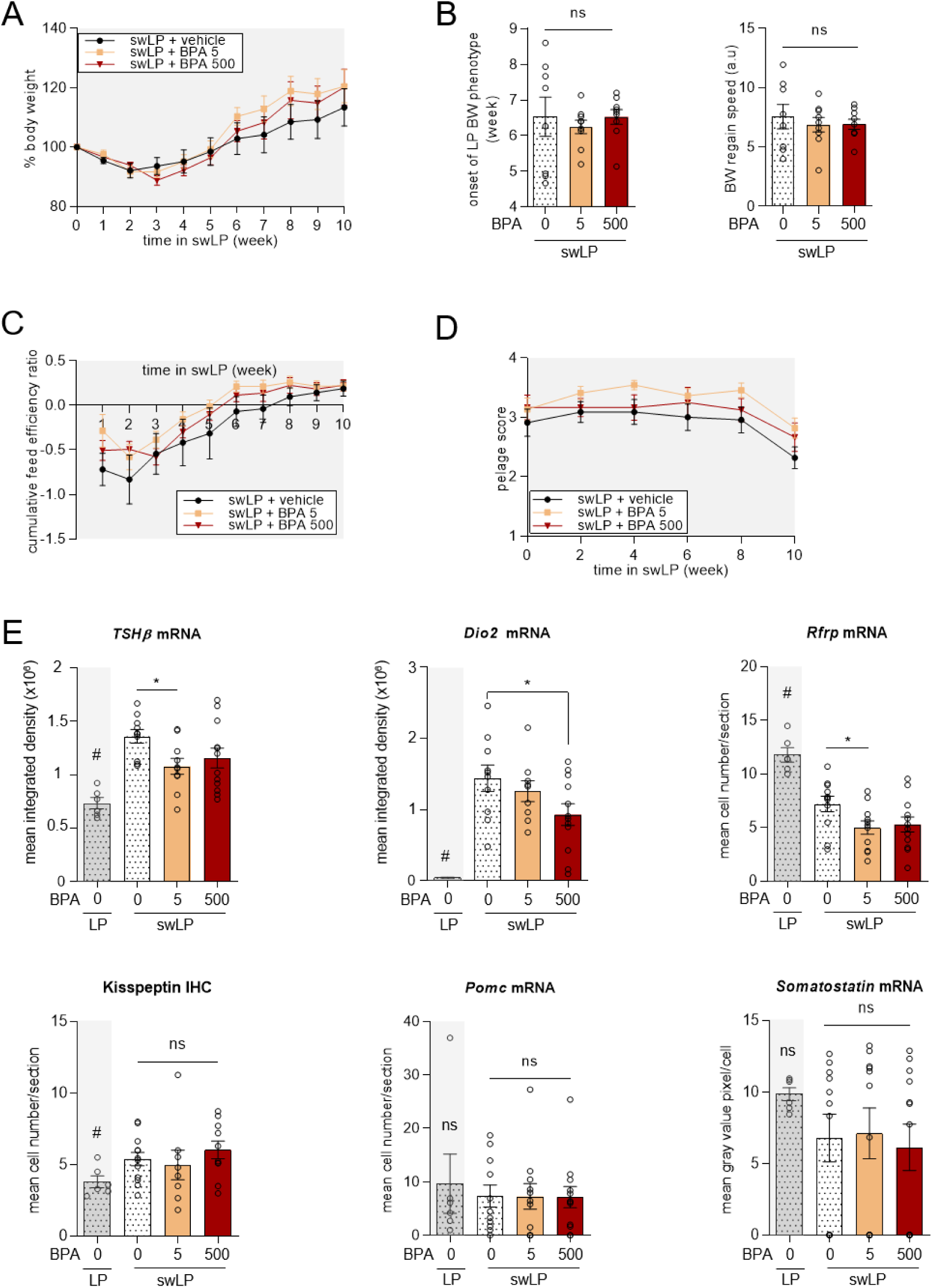
Effects of BPA exposure on body weight, food intake, pelage whitening, and various photoperiodic genes/proteins in male Djungarian hamsters transferred from short to long photoperiod. Male Djungarian hamsters adapted to short photoperiod (SP) were switched back in long photoperiod (swLP) and either fed with pellet food containing 0.02% ethanol (swLP + vehicle; n=11), or containing 5 (swLP + BPA 5; n=11) or 500 (swLP + BPA 500; n=12) µg BPA/kg/day. **(A)** Body weight (BW) expressed in percentage of the value before the swLP transfer. **(B)** Time in weeks to reach the LP body weight phenotype (point of inflection calculated from a non-linear regression model fitted to individual body weight raw data); and speed of LP-induced body weight gain (slope of the non-linear regression). **(C)** Cumulative feed efficiency ratio over 10 weeks of swLP. **(D)** Pelage score (from 1 being a LP-adapted grey color to 5 being a SP-adapted white color) over 10 weeks of swLP. **(E)** Expression of *pars tuberalis TSHβ*, and hypothalamic *Dio2*, *Rfrp*, *Pomc*, and *somatostatin* gene and kisspeptin after 10 weeks of swLP, or constant LP. Values are given as mean ± SEM (n=11 to 12 according to experimental groups). Statistical significance: for multiple comparisons between groups (B, E), *, p<0.05 vs swLP + vehicle; to ascertain the photoperiodic effect in gene/protein expression, #, p<0.05 between LP + vehicle and swLP + vehicle. *ns*, no significance.

After 10 weeks of swLP, male hamsters had regained high relative testis weights, as well as circulating testosterone concentrations (**Table 2**) characteristic of sexually active LP-adapted animals. BPA exposure did not modify these reproductive indexes.

Expression levels of the photoperiodic genes *TSHβ,* Dio2, and *Rfrp* 10 weeks after the switch from SP to LP did not reach values similar to those observed in hamsters kept constantly in LP, suggesting a still intermediate state at this stage (**Figure 6E**). Furthermore, BPA exposure during swLP reduced the expression of *TSHβ* and *Rfrp* (at the dose of 5 µg/kg/day) and *Dio2* (at the dose of 500 µg/kg/day) (**Figure 6E**). In contrast, the expression of the metabolic genes *Pomc* and *somatostatin,* and the reproductive protein kisspeptin, were not altered by either dose of BPA exposure after 10 weeks of swLP (**Figure 6E**).

## Discussion

The present study is, to the best of our knowledge, the first investigation of the impact of BPA exposure on the photoperiodic adaptation of a seasonal mammal. It demonstrates that BPA-enriched food disrupts the kinetics of Djungarian hamster’s physiological adaptation to seasonal changes in photoperiod, underscoring a marked sex-dependent effect of BPA on seasonal physiology.

### BPA exposure exerts a negligible effect on the integration of the photoperiodic message

Photoperiodic integration is known to involve a melatonin-driven change in *pars tuberalis* TSH action on hypothalamic thyroid hormone metabolism and RFRP-3 neurons (Dardente and Simonneaux, 2022 for review). In our study, none of the investigated doses of BPA (from 5 to 500 µg/kg/day) modified the SP-induced decrease in *TSHβ* and *Rfrp* mRNA and the increase in *Dio3* mRNA in both male and female hamsters. This suggests that BPA does not alter the SP melatonin signaling up to the RFRP-3 neurons. The LP-induced increase in *TSHβ, Dio2, and Rfrp* mRNA levels was not affected by BPA in female hamsters, but in males BPA exposure reduced *TSHβ, Dio2, and Rfrp* levels. This could result from BPA acting either on the melatoninergic message (potentially synthesis, transport, degradation, or MT1 receptors in the *pars tuberalis*) or directly on TSHβ synthesis. The effects of BPA on melatonin synthesis and action are not known, as the majority of experimental studies assessing the effects of BPA are based on melatonin-deficient mouse models or are not conducted in seasonal/photoperiodic contexts. Nevertheless, this BPA-induced reduction in the *TSHβ/Dio2/Rfrp* pathway in male hamsters was not associated with an obvious disruption of metabolic or reproductive responses to the LP transfer.

### Metabolic adaptation to photoperiodic changes is altered by BPA with marked sex-specific differences

BPA exposure significantly disturbed the kinetics of the Djungarian hamster’s metabolic adaptation to photoperiodic changes, revealing marked differences based on both the photoperiodic protocol and the hamster’s sex. Specifically, exposure to BPA during the LP to SP transfer slowed the acquisition of the lean metabolic phenotype in females exposed to 500 µg/kg/day but accelerated it in males exposed to 5 µg/kg/day. On the other hand, exposure to 5 or 500 µg/kg/day BPA during the SP to LP transfer accelerated the acquisition of the fat metabolic phenotype in females but had no obvious metabolic effect in males.

BPA has been previously identified as an obesogenic substance in multiple *in vitro* and *in vivo* studies (Moghaddam et al., 2015; Rubin et al., 2001; Angle et al., 2013; Marmugi et al., 2012; Alonso-Magdalena et al., 2006, 2010) and its exposure has been linked to increased risks of obesity and type 2 diabetes in humans (Lang et al., 2008; Sun et al., 2014; Song et al., 2016). The observation that female Djungarian hamsters exposed to BPA show a slowdown in body weight loss and a decrease in feed efficiency during the transfer to SP, as well as an acceleration in body weight gain and an increase in feed efficiency in swLP is therefore consistent with this obesogenic effect of BPA. The SP-induced decrease in Djungarian hamster’s body weight is known to depend on a TSH/T3-driven increase in ARC *somatostatin* expression (Dumbell et al., 2015; Klosen et al., 2013). Therefore, the slower rate of SP-induced body weight loss in female hamsters exposed to 500 µg/kg/day BPA could be attributed to the reduced expression of ARC *somatostatin* observed at 10 weeks of SP. These results are in line with previous studies reporting that BPA can alter the binding activities of the somatostatin subtype 3 receptors in the rat ARC (Facciolo et al., 2005), inhibit the synthesis and *in vitro* release of growth hormone (GH) via an alteration in the cellular signal transduction systems of GHRH (Katoh et al., 2004), and alter ERα receptors in a small population of ARC somatostatin neurons (Scanlan et al., 2003).

The photoperiodic regulation in female Djungarian hamster’s body weight is also associated with changes in *Pomc* gene expression, known to be higher in LP compared to SP (Cázarez-Márquez et al., 2019; Mercer et al., 2000; Reddy et al., 1999). Therefore, the accelerated body weight gain and improved feeding efficiency observed in female hamsters exposed to 5 µg/kg/day BPA during the SP to LP transfer may result from the observed reduction in the ARC *Pomc* expression. Previous studies have reported that BPA can disrupt *Pomc* expression, with contradictory variations depending on the study (Desai et al., 2018; Salehi et al., 2019) and induce a reduction in the number of POMC neuron projections to the paraventricular nuclei (Mackay et al., 2013). BPA effect on *Pomc* expression could also be mediated by a direct antagonistic disruption of T3, which has been shown to induce *Pomc* expression by binding to TRβ1 in POMC neurons (Bao et al., 2019).

In male Djungarian hamsters transferred from LP to SP, the acceleration of body weight loss and reduction in feed efficiency induced by exposure to 5 µg/kg/day BPA was unexpected given the recognized obesogenic effect of BPA. However, BPA exposure in humans is sometimes associated with a reduction in body mass indexes (Harley et al., 2013). No change in *Pomc* and *somatostatin* gene expression was associated with this BPA-induced metabolic alteration, but the glucose/insulin ratio was increased in male hamsters exposed to 5 µg/kg/day BPA as compared to control animals. This suggests an increase in insulin sensitivity and a reinforcement of the central anorexigenic effect of insulin (Kauffman and Castracane, 2003), as insulin can inhibit orexigenic NPY neurons (Lee and Herzog, 2021).

In order to better understand the metabolic properties of BPA in Djungarian hamsters, further investigations on the central regulation of metabolic neurons, notably their sensibility to leptin (Mackay et al., 2017), and on peripheral organs (liver, pancreas, adipose tissue, reported to be sensitive to BPA (Marmugi et al., 2012; Soriano et al., 2012; García-Arevalo et al., 2014) are required.

### BPA modulates the kinetics of SP-induced fur whitening in both female and male hamsters

Melatonin controls photoperiodic changes in fur color and density in seasonal rodents through the regulation of prolactin secretion (Hoffmann, 1973; Duncan et al., 1985; Rose et al., 1985). Exposure to 5 or 500 µg BPA/kg/day during the transfer from LP to SP resulted in a delay in females and an advance in males of fur whitening. Although technically challenging, the longitudinal analysis of the photoperiodic change in prolactin secretion could help to determine whether this sex-dependent effect of BPA occurs at the level of prolactin synthesis or on downstream mechanisms. One study in Agouti mice reported that BPA exposure induced a change in coat color mediated by a reduction in methylation of the Agouti gene in germ cells (Dolinoy et al., 2007). During the 10-week transfer from SP to LP, the fur color did not change significantly, independently of BPA exposure, possibly because the restoration of the LP grey fur phenotype takes more time.

### A low dose of BPA accelerates the LP-induced restoration of reproduction in female hamsters

A non-invasive and not stressful longitudinal monitoring of reproductive activity is challenging in the Djungarian hamster. Thus, the assessment of BPA’s effect on the kinetic of photoperiodic changes in reproductive activity could only be conducted in female hamsters transferred from SP to LP and sampled at an intermediate stage (5^th^ week of swLP). Exposure to 5 µg BPA/kg/day, but not 500 µg BPA/kg/day, accelerated uterine and ovarian weight gain compared to vehicle-treated female hamsters. Previous studies in birds have shown that other EDCs, specifically isoflavones, could also alter the dynamics of reproductive responses to changes in photoperiods. Notably, a diet enriched with isoflavones delayed the growth of the cloacal protuberance of oscines in response to an LP transfer (Corbitt et al., 2007) and reduced the testicular development of Japanese quails transferred from an inhibitory SP to a stimulatory LP (Wilhelms et al., 2006). To confirm that BPA exposure can alter the kinetics of the photoperiodic regulation of reproductive activity, complementary studies incorporating more intermediate sampling times should be conducted in male and female hamsters.

Recent studies in Djungarian hamsters have reported that the photoperiodic control of reproductive activity depends on ARC kisspeptin regulation by melatonin and sex steroids and that kisspeptin can restore SP-inhibited gonadal activity (Rasri-Klosen et al. 2017; Cázarez-Márquez et al., 2019). Interestingly, the early reactivation of the reproductive axis in females exposed to BPA during the LP transfer was associated with an advance in the reduction of ARC kisspeptin compared to control females. Previous studies have also reported that BPA can reduce kisspeptin ARC expression in rats (Patisaul et al., 2009) and mice (Ruiz-Pino et al., 2019), and suppress kisspeptin liberation in the median eminence of rhesus monkeys (Kurian et al., 2015). As ARC ERα expression can also be altered by BPA exposure (Cao et al., 2012; Monje et al., 2010) one hypothesis could be that BPA may induce an early sensitization of the ARC kisspeptin neurons to the negative feedback of circulating estrogens, which are already high at the 5^th^ week of swLP.

### Potential ecological impact of BPA exposure in Djungarian hamsters

Our study has reported clear BPA effects on the dynamics of photoperiodic adaptation in male and female Djungarian hamsters. From an ecological perspective, the acceleration of physiological adaptations to the winter photoperiod, as observed in BPA-exposed male hamsters, or to the summer photoperiod, as observed in BPA-exposed female hamsters, can be interpreted as an adaptive advantage, enabling the individuals to anticipate their physiology more quickly for the upcoming season. Nevertheless, animals may temporarily find themselves in a discordant physiological state with the seasonal change in their environment.

Our study also highlighted a striking sexual dimorphism in the BPA-induced change in the kinetics of photoperiodic integration, notably from LP to SP conditions where females showed a delay while males showed an advance in their metabolic adaptation. The consequences of such mismatches, even if temporary, between the physiological state of an individual, the resources of its environment, and the physiological state of its conspecifics may lead to impaired species adaptation to seasons at a population level.

## Conclusion

This study is the first to reveal significant effects of BPA exposure on the seasonal adaptation of a mammalian species, namely the Djungarian hamster. Findings demonstrate that BPA exposure disrupts the kinetics of metabolic and reproductive responses to seasonal changes in a sex-dependent manner. Seasonal adaptation relies on a complex photoneuroendocrine pathway involving the melatoninergic system, hypothalamic thyroid hormones and neuropeptides, and sex steroids, which are key targets for BPA and other EDCs. However, the effects of EDCs on seasonal physiology and its neuroendocrine mechanisms are still under-evaluated. Notably, the long-term effects of EDCs exposure on seasonal adaptation should be assessed, particularly when individuals are exposed during critical windows of vulnerability in early life. Additionnally, considering the ecological perspective, understanding the impact of EDCs on animals’ interaction within their ecosystem is essential for a comprehensive evaluation of the consequences of environmental contamination.

## Acknowledgments

This work was supported by the French agency for food, environmental and occupational health and safety (ANSES), the French Society for Endocrinology (SFE), and the Interdisciplinary Thematic Institute NeuroStra (ANR-10-IDEX-0002). We thank Dominique Ciocca, Sophie Reibel-Foisset, Nicolas Lethenet, and all the staff of the Chronobiotron’s animal facility of Strasbourg for taking care of the hamsters, as well as Paul Klosen for his guidance on the neuroanatomical analyses.

## Supplementary materials

**Table 1.** LC-MS/MS conditions for testosterone. LC and MS/MS conditions for the purification, detection and quantification of testosterone as well as their respective heavy-tagged counterparts. The flow rate was set at 90 µl/min on a ZORBAX SB-C18 column (150 x 1mm, 3.5μm) and the column was heated at 40°C.

| Mobile phase |  |  |  |
| --- | --- | --- | --- |
| Gradient | ACN | H<sub>2</sub>O | Formic acid |
| Mobile phase A | 1% | 98.9% | 0.1% |
| Mobile phase B | 99.9% | 0 | 0.1% |

| HPLC gradient |  |  |  |  |  |  |  |
| --- | --- | --- | --- | --- | --- | --- | --- |
| Time (min) | 0 | 2.5 | 5 | 11 | 13 | 13.5 | 20 |
| % B mobile phase | 0 | 0 | 20 | 98 | 98 | 0 | 0 |

| MS parameters |  |
| --- | --- |
| Mode | positive |
| Spray voltage | 3,500 V |
| Nebulizer gas | Nitrogen |
| Desolvation (nitrogen) sheath gas | 18 Arb |
| Aux gas | 7 Arb |
| Ion transfer tube temperature | 297°C |
| Vaporizer temperature | 131°C |
| Q1 and Q3 resolutions | 0.7 FWHM |
| Collision gas (CID, argon) pressure | 2 mTorr |

| Compound | Polarity | Precursor (m/z) | Product (m/z) | Ion product type | Collision Energy (V) | RF Lens (V) |
| --- | --- | --- | --- | --- | --- | --- |
| Testosterone-keto | Positive | 403.3 | 152.11 | Qualification | 40.64 | 158 |
| Testosterone-keto | Positive | 403.3 | 164.11 | Qualification | 41.29 | 158 |
| Testosterone-keto | Positive | 403.3 | 344.22 | Quantification | 20.41 | 158 |
| 2H4-testo-keto-407 | Positive | 407.32 | 153.11 | Qualification | 41.19 | 179 |
| 2H4-testo-keto-407 | Positive | 407.32 | 165.11 | Qualification | 42.76 | 179 |
| 2H4-testo-keto-407 | Positive | 407.32 | 348.22 | Quantification | 20.61 | 179 |

**Table 2.** LC-MS/MS conditions BPA-Glucuronide. LC and MS/MS conditions for the purification, detection and quantification of BPA-glucuronide as well as their respective heavy-tagged counterparts. The flow rate was set at 90 µl/min on a ZORBAX SB-C18 column (150 x 1mm, 3.5μm) and the column was heated at 40°C.

| HPLC gradient |  |  |  |  |  |  |  |
| --- | --- | --- | --- | --- | --- | --- | --- |
| Time (min) | 0 | 2.5 | 5 | 9 | 10.5 | 11.5 | 16 |
| % B mobile phase | 0 | 0 | 20 | 98 | 98 | 0 | 0 |

| MS parameters |  |
| --- | --- |
| Mode | Negative |
| Spray voltage | 2500 V |
| Nebulizer gas | Nitrogen |
| Desolvation (nitrogen) sheath gas | 18 Arb |
| **Aux gas** | 7 Arb |
| **Ion transfer tube temperature** | 297°C |
| **Vaporizer temperature** | 131°C |
| **Q1 and Q3 resolutions** | 0.7 FWHM |
| **Collision gas (CID, argon) pressure** | 2 mTorr |

MS ionization, selection, fragmentation and identification parameters
| **Compound** | **Polarity** | **Precursor (m/z)** | **Product (m/z)** | **Ion product type** | **Collision Energy (V)** | **RF Lens (V)** |
| --- | --- | --- | --- | --- | --- | --- |
| BPA-Glucuronide | Negative | 402.98 | 175.05 | Qualification | 15.20 | 185 |
| BPA-Glucuronide | Negative | 402.98 | 212.04 | Qualification | 37.40 | 185 |
| BPA-Glucuronide | Negative | 402.98 | 227.04 | Quantification | 26.28 | 185 |
| BPA-Glucuronide-<sup>13</sup>C12 | Negative | 415.35 | 174.98 | Qualification | 14.55 | 225 |
| BPA-Glucuronide-<sup>13</sup>C12 | Negative | 415.35 | 224.04 | Qualification | 36.74 | 225 |
| BPA-Glucuronide-<sup>13</sup>C12 | Negative | 415.35 | 239.02 | Quantification | 27.49 | 225 |

## References

1. Alonso-Magdalena, P., Morimoto, S., Ripoll, C., Fuentes, E., & Nadal, A. (2006). The estrogenic effect of bisphenol A disrupts pancreatic β-cell function in vivo and induces insulin resistance. Environmental health perspectives, 114(1), 106–112.

2. Alonso-Magdalena, P., Vieira, E., Soriano, S., Menes, L., Burks, D., Quesada, I., & Nadal, A. (2010). Bisphenol A exposure during pregnancy disrupts glucose homeostasis in mothers and adult male offspring. Environmental health perspectives, 118(9), 1243–1250.

3. Angle, B. M., Do, R. P., Ponzi, D., Stahlhut, R. W., Drury, B. E., Nagel, S. C., … Taylor, J. A. (2013). Metabolic disruption in male mice due to fetal exposure to low but not high doses of bisphenol A (BPA): evidence for effects on body weight, food intake, adipocytes, leptin, adiponectin, insulin and glucose regulation. Reprod Toxicol, 42, 256–268. doi:10.1016/j.reprotox.2013.07.017

4. Bankhead, P., Loughrey, M. B., Fernández, J. A., Dombrowski, Y., McArt, D. G., Dunne, P. D., … Coleman, H. G. (2017). QuPath: Open source software for digital pathology image analysis. Scientific Reports, 7(1), 1–7.

5. Bao, R., Onishi, K. G., Tolla, E., Ebling, F. J., Lewis, J. E., Anderson, R. L., … Stevenson, T. J. (2019). Genome sequencing and transcriptome analyses of the Siberian hamster hypothalamus identify mechanisms for seasonal energy balance. Proceedings of the National Academy of Sciences, 116(26), 13116–13121.

6. Bartness, T. J., & Wade, G. N. (1984). Photoperiodic control of body weight and energy metabolism in Syrian hamsters (Mesocricetus auratus): role of pineal gland, melatonin, gonads, and diet. Endocrinology, 114(2), 492–498.

7. Bockmann, J., Böckers, T. M., Vennemann, B., Niklowitz, P., Müller, J., Wittkowski, W., … Kreutz, M. R. (1996). Short photoperiod-dependent down-regulation of thyrotropin-alpha and –beta in hamster pars tuberalis-specific cells is prevented by pinealectomy. Endocrinology, 137(5), 1804–1813. doi:10.1210/endo.137.5.8612518

8. Bronson, F. H. (1985). Mammalian reproduction: an ecological perspective. Biol Reprod, 32(1), 1–26. doi:10.1095/biolreprod32.1.1

9. Butler, M. P., Turner, K. W., Park, J. H., Butler, J. P., Trumbull, J. J., Dunn, S. P., … Zucker, I. (2007). Simulated natural day lengths synchronize seasonal rhythms of asynchronously born male Siberian hamsters. Am J Physiol Regul Integr Comp Physiol, 293(1), R402–412. doi:10.1152/ajpregu.00146.2007

10. Cao, J., Mickens, J. A., McCaffrey, K. A., Leyrer, S. M., & Patisaul, H. B. (2012). Neonatal Bisphenol A exposure alters sexually dimorphic gene expression in the postnatal rat hypothalamus. Neurotoxicology, 33(1), 23–36. doi:10.1016/j.neuro.2011.11.002

11. Cázarez-Márquez, F., Milesi, S., Laran-Chich, M. P., Klosen, P., Kalsbeek, A., & Simonneaux, V. (2019). Kisspeptin and RFRP 3 modulate body mass in Phodopus sungorus via two different neuroendocrine pathways. Journal of neuroendocrinology, 31(4), e12710.

12. Corbitt, C., Satre, D., Adamson, L. A., Cobbs, G. A., & Bentley, G. E. (2007). Dietary phytoestrogens and photoperiodic response in a male songbird, the Dark-eyed Junco (Junco hyemalis). Gen Comp Endocrinol, 154(1-3), 16–21. doi:10.1016/j.ygcen.2007.06.026

13. Dardente, H., Menet, J. S., Poirel, V. J., Streicher, D., Gauer, F., Vivien-Roels, B., Masson-Pévet, M. (2003). Melatonin induces Cry1 expression in the pars tuberalis of the rat. Brain Res Mol Brain Res, 114(2), 101–106. doi:10.1016/s0169-328x(03)00134-7

14. Dardente, H., & Simonneaux, V. (2022). GnRH and the photoperiodic control of seasonal reproduction: Delegating the task to kisspeptin and RFRP-3. Journal of neuroendocrinology, 34(5), e13124.

15. Delfosse, V., Grimaldi, M., Pons, J. L., Boulahtouf, A., le Maire, A., Cavailles, V., Balaguer, P. (2012). Structural and mechanistic insights into bisphenols action provide guidelines for risk assessment and discovery of bisphenol A substitutes. Proc Natl Acad Sci U S A, 109(37), 14930–14935. doi:10.1073/pnas.1203574109

16. Desai, M., Ferrini, M. G., Han, G., Jellyman, J. K., & Ross, M. G. (2018). In vivo maternal and in vitro BPA exposure effects on hypothalamic neurogenesis and appetite regulators. Environmental research, 164, 45–52.

17. Dolinoy, D. C., Huang, D., & Jirtle, R. L. (2007). Maternal nutrient supplementation counteracts bisphenol A-induced DNA hypomethylation in early development. Proc Natl Acad Sci U S A, 104(32), 13056–13061. doi:10.1073/pnas.0703739104

18. Dumbell, R. A., Scherbarth, F., Diedrich, V., Schmid, H. A., Steinlechner, S., & Barrett, P. (2015). Somatostatin agonist pasireotide promotes a physiological state resembling short-day acclimation in the photoperiodic male Siberian hamster (Phodopus sungorus). Journal of neuroendocrinology, 27(7), 588–599.

19. Duncan, M. J., & Goldman, B. D. (1984). Hormonal regulation of the annual pelage color cycle in the Djungarian hamster, Phodopus sungorus. I. Role of the gonads and pituitary. J Exp Zool, 230(1), 89–95. doi:10.1002/jez.1402300112

20. Duncan, M. J., Goldman, B. D., Di Pinto, M. N., & Stetson, M. H. (1985). Testicular function and pelage color have different critical daylengths in the Djungarian hamster, Phodopus sungorus sungorus. Endocrinology, 116(1), 424–430. doi:10.1210/endo-116-1-424

21. Facciolo, R. M., Madeo, M., Alò, R., Canonaco, M., & Dessì-Fulgheri, F. (2005). Neurobiological effects of bisphenol A may be mediated by somatostatin subtype 3 receptors in some regions of the developing rat brain. Toxicol Sci, 88(2), 477–484. doi:10.1093/toxsci/kfi322

22. Figala, J., Hoffmann, K., & Goldau, G. (1973). [The annual cycle in the Djungarian Hamster Phodopus sungorus Pallas]. Oecologia, 12(2), 89–118. doi:10.1007/bf00345511

23. García-Arevalo, M., Alonso-Magdalena, P., Rebelo Dos Santos, J., Quesada, I., Carneiro, E. M., & Nadal, A. (2014). Exposure to bisphenol-A during pregnancy partially mimics the effects of a high-fat diet altering glucose homeostasis and gene expression in adult male mice. PloS one, 9(6), e100214.

24. Gorman, M. R., & Zucker, I. (1997). Environmental induction of photononresponsiveness in the Siberian hamster, Phodopus sungorus. Am J Physiol, 272(3 Pt 2), R887–895. doi:10.1152/ajpregu.1997.272.3.R887

25. Greives, T. J., Kriegsfeld, L. J., & Demas, G. E. (2008). Exogenous kisspeptin does not alter photoperiod-induced gonadal regression in Siberian hamsters (Phodopus sungorus). General and comparative endocrinology, 156(3), 552–558.

26. Guadaño-Ferraz, A., Obregón, M. J., Germain, D. L. S., & Bernal, J. (1997). The type 2 iodothyronine deiodinase is expressed primarily in glial cells in the neonatal rat brain. Proceedings of the National Academy of Sciences, 94(19), 10391–10396.

27. Hanon, E., Routledge, K., Dardente, H., Masson-Pévet, M., Morgan, P. J., & Hazlerigg, D. G. (2010). Effect of photoperiod on the thyroid-stimulating hormone neuroendocrine system in the European hamster (Cricetus cricetus). Journal of neuroendocrinology, 22(1), 51–55.

28. Harley, K. G., Gunier, R. B., Kogut, K., Johnson, C., Bradman, A., Calafat, A. M., & Eskenazi, B. (2013). Prenatal and early childhood bisphenol A concentrations and behavior in school-aged children. Environ Res, 126, 43–50. doi:10.1016/j.envres.2013.06.004

29. Hazlerigg, D., & Simonneaux, V. (2015). Seasonal regulation of reproduction in mammals. Knobil and Neill’s physiology of reproduction, 4.

30. Helfer, G., Ross, A., & Morgan, P. (2013). Neuromedin U Partly Mimics Thyroid-Stimulating Hormone and Triggers W nt/β-Catenin Signalling in the Photoperiodic Response of F 344 Rats. Journal of neuroendocrinology, 25(12), 1264–1272.

31. Henson, J. R., Carter, S. N., & Freeman, D. A. (2013). Exogenous T3 elicits long day–like alterations in testis size and the RFamides kisspeptin and gonadotropin-inhibitory hormone in short-day Siberian hamsters. Journal of biological rhythms, 28(3), 193–200.

32. Herwig, A., de Vries, E. M., Bolborea, M., Wilson, D., Mercer, J. G., Ebling, F. J., … Barrett, P. (2013). Hypothalamic ventricular ependymal thyroid hormone deiodinases are an important element of circannual timing in the Siberian hamster (Phodopus sungorus). PloS one, 8(4), e62003.

33. Herwig, A., Petri, I., & Barrett, P. (2012). Hypothalamic gene expression rapidly changes in response to photoperiod in juvenile Siberian hamsters (Phodopus sungorus). J Neuroendocrinol, 24(7), 991–998. doi:10.1111/j.1365-2826.2012.02324.x

34. Hoffmann, K. (1973). The influence of photoperiod and melatonin on testis size, body weight, and pelage colour in the Djungarian hamster (Phodopus sungorus). Journal of comparative physiology, 85(3), 267–282. doi:10.1007/BF00694233

35. Jasnow, A. M., Huhman, K. L., Bartness, T. J., & Demas, G. E. (2000). Short-day increases in aggression are inversely related to circulating testosterone concentrations in male Siberian hamsters (Phodopus sungorus). Horm Behav, 38(2), 102–110. doi:10.1006/hbeh.2000.1604

36. Katoh, K., Matsuda, A., Ishigami, A., Yonekura, S., Ishiwata, H., Chen, C., & Obara, Y. (2004). Suppressing effects of bisphenol A on the secretory function of ovine anterior pituitary cells. Cell Biol Int, 28(6), 463–469. doi:10.1016/j.cellbi.2004.03.016

37. Kauffman, R. P., & Castracane, V. D. (2003). Assessing insulin sensitivity.(Controlling PCOS, part 1). Contemporary OB/GYN, 48(1), 30–39.

38. Kitamura, S., Suzuki, T., Sanoh, S., Kohta, R., Jinno, N., Sugihara, K., … Ohta, S. (2005). Comparative study of the endocrine-disrupting activity of bisphenol A and 19 related compounds. Toxicol Sci, 84(2), 249–259. doi:10.1093/toxsci/kfi074

39. Klosen, P., Bienvenu, C., Demarteau, O., Dardente, H., Guerrero, H., Pévet, P., & Masson-Pévet, M. (2002). The mt1 melatonin receptor and RORbeta receptor are co-localized in specific TSH-immunoreactive cells in the pars tuberalis of the rat pituitary. J Histochem Cytochem, 50(12), 1647–1657. doi:10.1177/002215540205001209

40. Klosen, P., Maessen, X., & van den Bosch de Aguilar, P. (1993). PEG embedding for immunocytochemistry: application to the analysis of immunoreactivity loss during histological processing. J Histochem Cytochem, 41(3), 455–463. doi:10.1177/41.3.8429209

41. Klosen, P., Sébert, M. E., Rasri, K., Laran-Chich, M. P., & Simonneaux, V. (2013). TSH restores a summer phenotype in photoinhibited mammals via the RF-amides RFRP3 and kisspeptin. The FASEB Journal, 27(7), 2677–2686.

42. Kuiper, G. G., Lemmen, J. G., Carlsson, B., Corton, J. C., Safe, S. H., van der Saag, P. T., … Gustafsson, J. A. (1998). Interaction of estrogenic chemicals and phytoestrogens with estrogen receptor beta. Endocrinology, 139(10), 4252–4263. doi:10.1210/endo.139.10.6216

43. Kurian, J. R., Keen, K. L., Kenealy, B. P., Garcia, J. P., Hedman, C. J., & Terasawa, E. (2015). Acute Influences of Bisphenol A Exposure on Hypothalamic Release of Gonadotropin-Releasing Hormone and Kisspeptin in Female Rhesus Monkeys. Endocrinology, 156(7), 2563–2570. doi:10.1210/en.2014-1634

44. Lang, I. A., Galloway, T. S., Scarlett, A., Henley, W. E., Depledge, M., Wallace, R. B., & Melzer, D. (2008). Association of urinary bisphenol A concentration with medical disorders and laboratory abnormalities in adults. Jama, 300(11), 1303–1310. doi:10.1001/jama.300.11.1303

45. Lee, N. J., & Herzog, H. (2021). Coordinated regulation of energy and glucose homeostasis by insulin and the NPY system. Journal of neuroendocrinology, 33(4), e12925.

46. Lopez-Rodriguez, D., Franssen, D., Bakker, J., Lomniczi, A., & Parent, A. S. (2021). Cellular and molecular features of EDC exposure: consequences for the GnRH network. Nat Rev Endocrinol, 17(2), 83–96. doi:10.1038/s41574-020-00436-3

47. Mackay, H., Patterson, Z. R., & Abizaid, A. (2017). Perinatal Exposure to Low-Dose Bisphenol-A Disrupts the Structural and Functional Development of the Hypothalamic Feeding Circuitry. Endocrinology, 158(4), 768–777. doi:10.1210/en.2016-1718

48. Mackay, H., Patterson, Z. R., Khazall, R., Patel, S., Tsirlin, D., & Abizaid, A. (2013). Organizational effects of perinatal exposure to bisphenol-A and diethylstilbestrol on arcuate nucleus circuitry controlling food intake and energy expenditure in male and female CD-1 mice. Endocrinology, 154(4), 1465–1475. doi:10.1210/en.2012-2044

49. Marmugi, A., Ducheix, S., Lasserre, F., Polizzi, A., Paris, A., Priymenko, N., … Martin, P. G. (2012). Low doses of bisphenol A induce gene expression related to lipid synthesis and trigger triglyceride accumulation in adult mouse liver. Hepatology, 55(2), 395–407.

50. Mercer, J. G., Moar, K. M., Logie, T. J., Findlay, P. A., Adam, C. L., & Morgan, P. J. (2001). Seasonally Inappropriate Body Weight Induced by Food Restriction: Effect on Hypothalamic Gene Expression in Male Siberian Hamsters. Endocrinology, 142(10), 4173–4181. doi:10.1210/endo.142.10.8454

51. Mercer, J. G., Moar, K. M., Ross, A. W., Hoggard, N., & Morgan, P. J. (2000). Photoperiod regulates arcuate nucleus POMC, AGRP, and leptin receptor mRNA in Siberian hamster hypothalamus. *American Journal of Physiology-Regulatory*, Integrative and Comparative Physiology, 278(1), R271–R281.

52. Mikkelsen, J. D., & Simonneaux, V. (2009). The neuroanatomy of the kisspeptin system in the mammalian brain. Peptides, 30(1), 26–33. doi:10.1016/j.peptides.2008.09.004

53. Milesi, S., Simonneaux, V., & Klosen, P. (2017). Downregulation of Deiodinase 3 is the earliest event in photoperiodic and photorefractory activation of the gonadotropic axis in seasonal hamsters. Scientific Reports, 7(1), 1–10.

54. Moghaddam, H. S., Samarghandian, S., & Farkhondeh, T. (2015). Effect of bisphenol A on blood glucose, lipid profile and oxidative stress indices in adult male mice. Toxicology mechanisms and methods, 25(7), 507–513.

55. Monje, L., Varayoud, J., Muñoz-de-Toro, M., Luque, E. H., & Ramos, J. G. (2010). Exposure of neonatal female rats to bisphenol A disrupts hypothalamic LHRH pre-mRNA processing and estrogen receptor alpha expression in nuclei controlling estrous cyclicity. Reprod Toxicol, 30(4), 625–634. doi:10.1016/j.reprotox.2010.08.004

56. Moralia, M.-A., Quignon, C., Simonneaux, M., & Simonneaux, V. (2022). Environmental disruption of reproductive rhythms. Frontiers in neuroendocrinology, 66, 100990.

57. Moriyama, K., Tagami, T., Akamizu, T., Usui, T., Saijo, M., Kanamoto, N., … Nakao, K. (2002). Thyroid hormone action is disrupted by bisphenol A as an antagonist. J Clin Endocrinol Metab, 87(11), 5185–5190. doi:10.1210/jc.2002-020209

58. Nakane, Y., & Yoshimura, T. (2019). Photoperiodic Regulation of Reproduction in Vertebrates. Annu Rev Anim Biosci, 7, 173–194. doi:10.1146/annurev-animal-020518-115216

59. Norris, D. O., & Jones, R. E. (1987). Hormones and reproduction in fishes, amphibians, and reptiles: Springer.

60. Patisaul, H. B., Todd, K. L., Mickens, J. A., & Adewale, H. B. (2009). Impact of neonatal exposure to the ERalpha agonist PPT, bisphenol-A or phytoestrogens on hypothalamic kisspeptin fiber density in male and female rats. Neurotoxicology, 30(3), 350–357. doi:10.1016/j.neuro.2009.02.010

61. Petri, I., Dumbell, R., Scherbarth, F., Steinlechner, S., & Barrett, P. (2014). Effect of exercise on photoperiod-regulated hypothalamic gene expression and peripheral hormones in the seasonal Dwarf Hamster Phodopus sungorus. PloS one, 9(3), e90253.

62. Rasri-Klosen, K., Simonneaux, V., & Klosen, P. (2017). Differential response patterns of kisspeptin and RFamide-related peptide to photoperiod and sex steroid feedback in the Djungarian hamster (Phodopus sungorus). Journal of neuroendocrinology, 29(9), e12529.

63. Reddy, A. B., Cronin, A. S., Ford, H., & Ebling, F. J. (1999). Seasonal regulation of food intake and body weight in the male Siberian hamster: studies of hypothalamic orexin (hypocretin), neuropeptide Y (NPY) andpro-opiomelanocortin (POMC). European Journal of Neuroscience, 11(9), 3255–3264.

64. Revel, F. G., Saboureau, M., Masson-Pévet, M., Pévet, P., Mikkelsen, J. D., & Simonneaux, V. (2006). Kisspeptin mediates the photoperiodic control of reproduction in hamsters. Current Biology, 16(17), 1730–1735.

65. Rose, J., Stormshak, F., Oldfield, J., & Adair, J. (1985). The effects of photoperiod and melatonin on serum prolactin levels of mink during the autumn molt. J Pineal Res, 2(1), 13–19. doi:10.1111/j.1600-079x.1985.tb00624.x

66. Rousseau, K., Atcha, Z., Cagampang, F. R. A., Le Rouzic, P., Stirland, J. A., Ivanov, T. R., … Loudon, A. S. (2002). Photoperiodic regulation of leptin resistance in the seasonally breeding Siberian hamster (Phodopus sungorus). Endocrinology, 143(8), 3083–3095.

67. Rubin, B. S., Murray, M. K., Damassa, D. A., King, J. C., & Soto, A. M. (2001). Perinatal exposure to low doses of bisphenol A affects body weight, patterns of estrous cyclicity, and plasma LH levels. Environmental health perspectives, 109(7), 675–680.

68. Ruiz-Pino, F., Miceli, D., Franssen, D., Vazquez, M. J., Farinetti, A., Castellano, J. M., … Tena-Sempere, M. (2019). Environmentally Relevant Perinatal Exposures to Bisphenol A Disrupt Postnatal Kiss1/NKB Neuronal Maturation and Puberty Onset in Female Mice. Environ Health Perspect, 127(10), 107011. doi:10.1289/ehp5570

69. Sáenz de Miera, C., Hanon, E. A., Dardente, H., Birnie, M., Simonneaux, V., Lincoln, G. A., & Hazlerigg, D. G. (2013). Circannual variation in thyroid hormone deiodinases in a short-day breeder. Journal of neuroendocrinology, 25(4), 412–421.

70. Salehi, A., Loganathan, N., & Belsham, D. D. (2019). Bisphenol A induces Pomc gene expression through neuroinflammatory and PPARγ nuclear receptor-mediated mechanisms in POMC-expressing hypothalamic neuronal models. Mol Cell Endocrinol, 479, 12–19. doi:10.1016/j.mce.2018.08.009

71. Scanlan, N., Dufourny, L., & Skinner, D. C. (2003). Somatostatin-14 neurons in the ovine hypothalamus: colocalization with estrogen receptor alpha and somatostatin-28(1-12) immunoreactivity, and activation in response to estradiol. Biol Reprod, 69(4), 1318–1324. doi:10.1095/biolreprod.103.017848

72. Schmidt, U., Weigert, M., Broaddus, C., & Myers, G. (2018). Cell detection with star-convex polygons. Paper presented at the Medical Image Computing and Computer Assisted Intervention–MICCAI 2018: 21st International Conference, Granada, Spain, September 16-20, 2018, Proceedings, Part II 11.

73. Shinomiya, A., Shimmura, T., Nishiwaki-Ohkawa, T., & Yoshimura, T. (2014). Regulation of seasonal reproduction by hypothalamic activation of thyroid hormone. Front Endocrinol (Lausanne), 5, 12. doi:10.3389/fendo.2014.00012

74. Song, Y., Chou, E. L., Baecker, A., You, N. C., Song, Y., Sun, Q., & Liu, S. (2016). Endocrine-disrupting chemicals, risk of type 2 diabetes, and diabetes-related metabolic traits: A systematic review and meta-analysis. J Diabetes, 8(4), 516–532. doi:10.1111/1753-0407.12325

75. Soriano, S., Alonso-Magdalena, P., Garcia-Arevalo, M., Novials, A., Muhammed, S. J., Salehi, A., … Nadal, A. (2012). Rapid insulinotropic action of low doses of bisphenol-A on mouse and human islets of Langerhans: role of estrogen receptor β. PloS one, 7(2), e31109.

76. Sun, Q., Cornelis, M. C., Townsend, M. K., Tobias, D. K., Eliassen, A. H., Franke, A. A., … Hu, F. B. (2014). Association of urinary concentrations of bisphenol A and phthalate metabolites with risk of type 2 diabetes: a prospective investigation in the Nurses’ Health Study (NHS) and NHSII cohorts. Environ Health Perspect, 122(6), 616–623. doi:10.1289/ehp.1307201

77. Tu, H. M., Kim, S.-W., Salvatore, D., Bartha, T., Legradi, G., Larsen, P. R., & Lechan, R. M. (1997). Regional distribution of type 2 thyroxine deiodinase messenger ribonucleic acid in rat hypothalamus and pituitary and its regulation by thyroid hormone. Endocrinology, 138(8), 3359–3368.

78. Wang, H., Ding, Z., Shi, Q. M., Ge, X., Wang, H. X., Li, M. X., … Xu, L. C. (2017). Anti-androgenic mechanisms of Bisphenol A involve androgen receptor signaling pathway. Toxicology, 387, 10–16. doi:10.1016/j.tox.2017.06.007

79. Warner, A., Jethwa, P. H., Wyse, C. A., I’anson, H., Brameld, J. M., & Ebling, F. J. (2010). Effects of photoperiod on daily locomotor activity, energy expenditure, and feeding behavior in a seasonal mammal. *American Journal of Physiology-Regulatory*, Integrative and Comparative Physiology, 298(5), R1409–R1416.

80. Watanabe, M., Yasuo, S., Watanabe, T., Yamamura, T., Nakao, N., Ebihara, S., & Yoshimura, T. (2004). Photoperiodic regulation of type 2 deiodinase gene in Djungarian hamster: possible homologies between avian and mammalian photoperiodic regulation of reproduction. Endocrinology, 145(4), 1546–1549.

81. Wilhelms, K. W., Scanes, C. G., & Anderson, L. L. (2006). Lack of estrogenic or antiestrogenic actions of soy isoflavones in an avian model: the Japanese quail. Poult Sci, 85(11), 1885–1889. doi:10.1093/ps/85.11.1885

82. Yoshimura, T., Yasuo, S., Watanabe, M., Iigo, M., Yamamura, T., Hirunagi, K., & Ebihara, S. (2003). Light-induced hormone conversion of T4 to T3 regulates photoperiodic response of gonads in birds. Nature, 426(6963), 178–181.

